# Multivalent interactions make adherens junction-cytoskeletal linkage robust during morphogenesis

**DOI:** 10.1101/2021.05.04.442628

**Authors:** Kia Z. Perez-Vale, Kristi D. Yow, Amy E. Byrnes, Tara M. Finegan, Kevin C. Slep, Mark Peifer

## Abstract

Embryogenesis requires cells to change shape and move without disrupting epithelial integrity. This requires robust, responsive linkage between adherens junctions and the actomyosin cytoskeleton. Using *Drosophila* morphogenesis we define molecular mechanisms mediating junction-cytoskeletal linkage and explore the role of mechanosensing. We focus on the junction-cytoskeletal linker Canoe, a multidomain protein. We engineered the *canoe* locus to define how its domains mediate its mechanism of action. To our surprise, the PDZ and FAB domains, which we thought connected junctions and F-actin, are not required for viability or mechanosensitive recruitment to junctions under tension. The FAB domain stabilizes junctions experiencing elevated force, but in its absence most cells recover, suggesting redundant interactions. In contrast, the Rap1-binding RA domains are critical for all Cno functions and for enrichment at junctions under tension. This supports a model in which junctional robustness derives from a large protein network assembled via multivalent interactions, with proteins at network nodes and some node connections more critical than others.

## Introduction

Epithelia are the principal animal tissue type, acting as barriers separating body compartments and lining body cavities. Their tissue architecture requires cells to adhere to one another and to the underlying extracellular matrix, and polarize by sending different proteins to apical and basal domains. However, epithelia are not static. During embryogenesis and tissue homeostasis cells must change shape and move while maintaining tissue integrity (Gillard and Roper, 2020). This requires force generation exerted by the actomyosin cytoskeleton linked to cell-cell and cell-matrix junctions. In adherens junctions (AJs), cell-cell adhesion is mediated by E-cadherin (Ecad), with Nectins also linking cells in some tissues.

The classic view of AJ-cytoskeletal linkage involves a simple linear model, in which homophilic Ecad interactions drive adhesion, and Ecad’s cytoplasmic tail binds beta-catenin (*β*cat), which in turn binds alpha- catenin (*α*cat), which then directly binds actin. However, this connectivity model is substantially oversimplified (Charras and Yap, 2018; Pinheiro and Bellaiche, 2018). First, AJ-cytoskeletal linkage is mechanosensitive, with force elevating *α*cat affinity for actin (Buckley et al., 2014; Yonemura et al., 2010). Second, many other linker proteins localize to AJs, including vinculin, Afadin/Canoe (Cno), ZO-1/Polychaetoid (Pyd), and Ajuba (Perez-Vale and Peifer, 2020). These observations motivated us to explore how this protein network permits cells to change shape and move while maintaining epithelial integrity.

*Drosophila* embryogenesis provides an outstanding place to study the machinery regulating morphogenesis, with many events requiring AJ-cytoskeletal linkage (Perez-Vale and Peifer, 2020). This begins at cellularization, when AJs assemble apically. During gastrulation, apical constriction drives mesoderm invagination and cell intercalation drives germband extension. Cells then remodel AJs to accommodate neighbors rounding up to divide or delaminating as neural stem cells, while collective cell migration drives epidermal dorsal closure. Often, force generation is planar polarized. For example, during germband extension myosin enrichment and force generation are highest at anterior-posterior (AP) borders and tricellular junctions (TCJs, where three or more cells meet). To maintain epithelial integrity, AJ-cytoskeletal linkages are modulated to reinforce junctions under elevated tension.

This led us to focus on Cno, Afadin’s fly homolog (Kuriyama et al., 1996; Miyamoto et al., 1995). Cno plays important roles during epithelial polarity establishment (Choi et al., 2013), mesoderm invagination (Sawyer et al., 2009), germband extension (Sawyer et al., 2011), dorsal closure (Boettner et al., 2003), and neuroblast division (Speicher et al., 2008). In Cno’s absence the cytoskeleton detaches from AJs, revealing an important linker role. Intriguingly, during germband extension, Cno is enriched at AP borders and TCJs, sites where force generation is highest (Fernandez-Gonzalez et al., 2009; Yu and Zallen, 2020). These sites are also where, in Cno’s absence, myosin detaches from AJs and AJ integrity weakened (Sawyer et al., 2009; 2011), though most cells in *cno* mutants ultimately recover epithelial architecture (Manning et al., 2019). Studies of gastrulating mouse embryos and MDCK cells revealed that Afadin plays similar roles in vertebrates (Ikeda et al., 1999; Zhadanov et al., 1999; Choi et al., 2016). In MDCK cells, Afadin knockdown disrupts AJ-cytoskeletal linkage, with the most pronounced effects at TCJs, where molecular tension on AJs is highest. However, Cno/Afadin does not act alone—a network of proteins act together with or in parallel with Cno, including Pyd/ZO-1 (Choi et al., 2011), Ajuba (Rauskolb et al., 2019; Razzell et al., 2018), Sidekick (Sdk; Finegan et al., 2019), and Smallish/LMO7 (Beati et al., 2018). Given Cno’s central place in this network, defining molecular mechanisms by which Cno is regulated, and by which it links AJs and the cytoskeleton to maintain AJ homoeostasis will move the field forward.

Cno is a multidomain scaffolding protein, in which some domains have known binding partners and others are less well characterized. We seek to define the roles of each domain in Cno’s mechanism of action. At the N-terminus are two Ras-association (RA) domains, which bind GTP-bound active Rap1(Boettner et al., 2003).

Rap1 activates and uses Cno as an effector (Kooistra et al., 2007). The RA domain is the only domain we previously manipulated, and in this case we only examined the earliest of Cno’s many roles, that in polarity establishment during cellularization. During cellularization Rap1 regulates Cno cortical recruitment (Sawyer et al., 2009). However, surprisingly, Cno’s RA domains are only important for some aspects of Cno localization at this stage, suggesting Rap1 regulates Cno via both RA-dependent and independent means (Bonello et al., 2018). Thus the RA domains are important for Cno’s role in initial apical-basal polarity establishment, but their roles later in morphogenesis remain untested.

Cno’s central region contains tandem Forkhead association (FHA), Dilute (DIL), and <u>P</u>SD95 <u>D</u>isc large <u>Z</u>O- 1 (PDZ) domains. While FHA and DIL binding partners/functions remain unknown, the PDZ binds the C-termini of Ecad (Sawyer et al., 2009) and nectins (Fujiwara et al., 2015; Takahashi et al., 1999; Wei et al., 2005), providing a connection to AJs. C-terminal to the PDZ, Cno and Afadin both have less well conserved intrinsically disordered linkers, in which binding sites for other proteins are embedded, including *α*cat (Sakakibara et al., 2020) and LGN (Carminati et al., 2016). Cno localization during polarity establishment also depends on filamentous actin (F-actin; Sawyer et al., 2009). While this is thought to be mediated by the C-terminal F-actin binding (FAB) domain (Mandai et al., 1997; Sawyer et al., 2009), other evidence suggests Cno cortical recruitment occurs via multivalent interactions (Yu and Zallen, 2020; Bonello et al., 2018). We now need to define the function of individual domains in Cno’s diverse morphogenetic functions.

The known roles of the PDZ, binding Ecad and nectins, and of the FAB, binding actin, suggest the hypothesis that Cno links AJs and the cytoskeleton through its PDZ and FAB domains (Sawyer et al., 2009). Alternately, each may be part of a multivalent set of connections. To distinguish these hypotheses, we used CRISPR/Cas9 to introduce into the locus site-directed mutants deleting individual domains. These data revealed surprising insights into roles of the PDZ, FAB, and RA domains, significantly revising Cno’s proposed mechanism of action.

## Results

### A platform to replace wildtype Cno with site-directed mutants

To define Cno’s mechanism of action, we examined how it uses its diverse protein interaction domains to link AJs and the cytoskeleton. The simplest hypothesis was that the PDZ provided the critical link to AJs, binding Ecad (Sawyer et al., 2009) and Echinoid (Ed; Wei et al., 2005), and the C-terminal FAB completed the link to actin (Mandai et al., 1997; Sawyer et al., 2009). We thus set out to generate mutants precisely deleting each domain. *cno* presented a challenge, with 21 exons over ∼50kbp (Fig 1A, Fig S1A). The first exon contains no protein coding sequence, and is separated by >13kbp from the second exon containing the start codon. We used CRISPR/Cas9 to delete exons 2-6, along with short portions of the flanking introns, replacing them with an attP site and selectable markers (Fig 1A). This allows subsequent integration of WT or mutant *cno* constructs via phiC31 integrase (Bischof et al., 2007).

**Fig 1.**
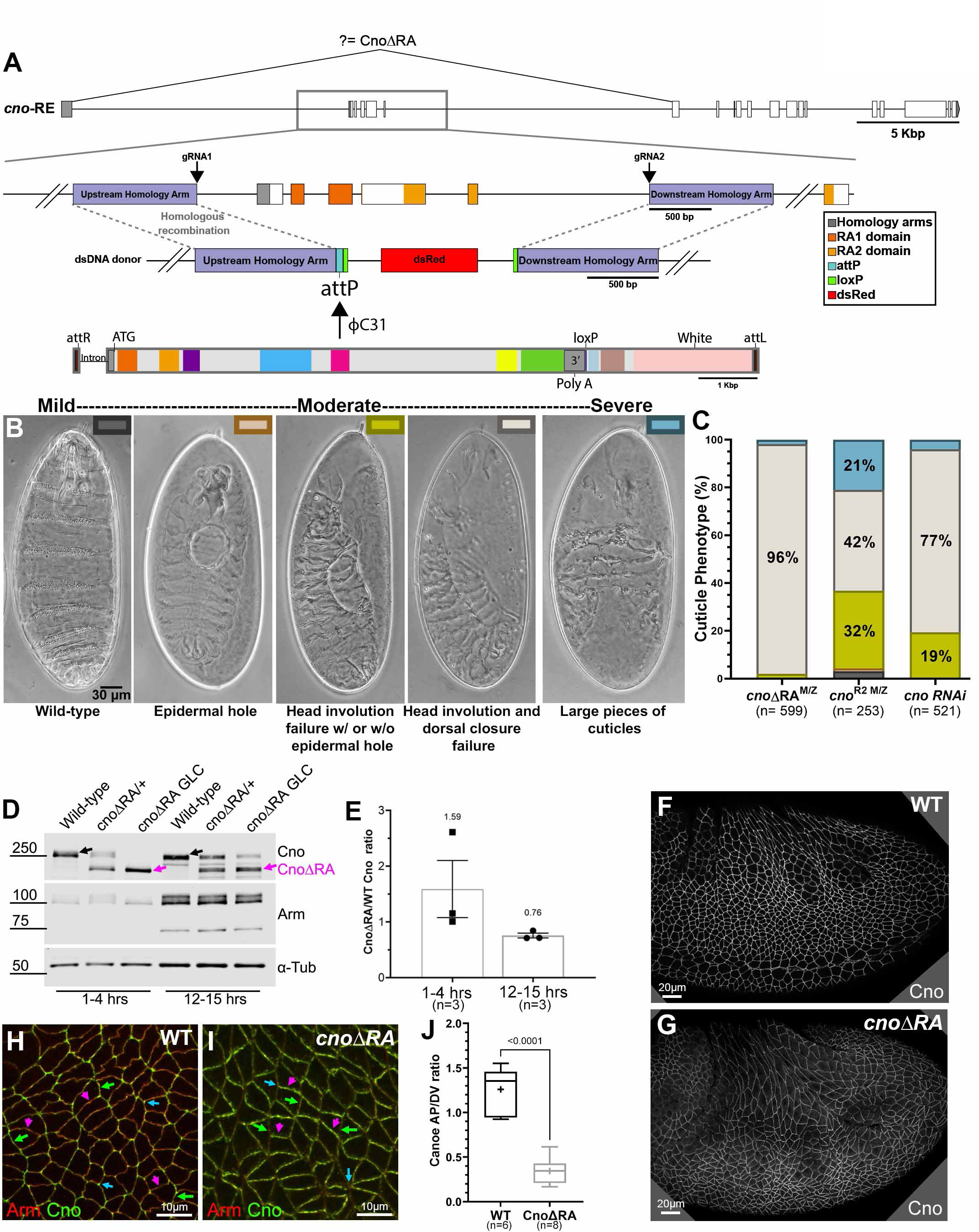
A platform for site-directed mutants and new insights into RA function. A. *cno* locus. Top: Exons and introns, Cno-PE isoform. Middle: Our CRISPR-generated deletion removes exons 2-6 (including start codon) and inserts an attP site and selectable marker (dsRed). Bottom. WT rescue construct. B. Representative cuticles of *cno* mutants. C. Deleting the RA domains substantially reduces Cno function in morphogenesis. D. Our new mutant produces WT levels of an N-terminally truncated Cno protein. Immunoblot, embryonic extracts, genotypes and times indicated (GLC=germline clone). *α*-tubulin = loading control. Black arrow=WT Cno. Magenta arrow= CnoΔRA. E. CnoΔRA protein levels relative to WT. F,G. CnoΔRA remains localized to AJs.H. WT Cno is enriched at TCJs (blue arrows) and at AP borders (green arrows). DV borders (magenta arrows). I. CnoΔRA is strongly enriched at DV borders (magenta vs green arrows). TCJ enrichment is lost (blue arrows). J. CnoΔRA planar polarity.

### This deletion led to a strong loss of function allele encoding a protein lacking the RA domains

Having engineered this deletion as the background for site-directed *cno* mutants, we examined its phenotype. As expected, it was homozygous lethal and lethal when heterozygous with our null allele, *cno^R2^*, which has an early stop codon and does not produce detectable protein (Sawyer et al., 2009). As we describe in detail below, viability was fully restored by integrating a WT GFP-tagged rescue construct. To define the effect of our new deletion on morphogenesis, we generated maternal/zygotic (M/Z) mutants, using the FLP/FRT approach (Chou and Perrimon, 1996; this and subsequent crosses are in Fig S2). 58% of progeny died, consistent with full M/Z lethality (50% of embryos) and some lethality of embryos receiving a paternal WT gene. As a first assessment of effects, we evaluated larval cuticles, allowing visualization of morphogenetic events and epithelial integrity (Fig 1B). M/Z mutants had a uniform phenotype: 96% had complete failure of both head involution and dorsal closure (Fig 1B,C). This is slightly more severe than the phenotype of embryos in which Cno was knocked down via RNAi to nearly undetectable levels (<5%; (Bonello et al., 2018), and was similar to that of *cno^R2^* M/Z mutants (from this point referred to as *cnoM/Z* null), though the fraction of embryos with severe defects in epidermal integrity was lower in our new mutant (2% vs 21%). We had a second indication that our new mutant was not null—while *cno^R2^* is zygotically homozygous embryonic lethal, albeit with only mild effects on morphogenesis (Sawyer et al., 2009), our new mutants died after embryogenesis.

Since we deleted the start codon and five exons this small amount of residual function surprised us. We thus examined whether this allele produced protein, immunoblotting samples from heterozygous and M/Z mutants. To our surprise, our new allele produced WT levels of a truncated protein (∼200 kDa; WT Cno is ∼250kDa) recognized by our anti-Cno antibody, raised to the C-terminal FAB (Fig 1D; 1.6 x WT at 1-4 hr; 0.76 x WT at 12-15 hr; Fig 1E). We suspect aberrant splicing joins exon 1 to exon 7 (Fig 1A), resulting in a protein with an alternative start codon. The first methionine in exon 7 is at amino acid 340, predicting a 189kD protein, roughly consistent with the protein observed. This protein lacks RA1 and most of RA2 (both removed by the deletion), and thus we refer to it as CnoΔRA. Intriguingly, CnoΔRA still localized to AJs (Fig 1F vs. G), as had a UAS-driven CnoΔRA protein overexpressed in a *cno* knockdown background (Bonello et al., 2018). Thus, our approach fortuitously generated a new allele allowing us to test roles of the RA domains. The cuticle analysis above suggested the RA domains are critical for morphogenesis.

### The RA domains are important for all Cno functions in morphogenesis, helping maintain epithelial integrity at junctions under tension

Rap1 is a key Cno regulator (Kooistra et al., 2007). Our new allele allowed us to define RA domain roles in morphogenesis at the cell biological level, by comparing *cnoΔRA* M/Z and *cnoM/Z* null mutants (Sawyer et al., 2009), our baseline for complete loss-of-function. As we describe below, *cnoΔRA* had many hallmark features of *cno* null mutants, suggesting the RA domains are critical for all Cno functions.

We first examined Cno’s role in apical-basal polarity establishment, when it positions AJ proteins and their partner Bazooka (Baz=Par3) apically. CnoΔRA remained cortical and roughly enriched apically (Fig S3A-C vs D-F), but some protein moved basally (Fig S3C vs F, yellow arrows), and TCJ cable enrichment was lost (Fig S3B’ vs E’, arrows; S3C vs F, green arrows). CnoΔRA localization to AJs was restored during gastrulation (Fig S3G vs H). In *cnoΔRA* mutants initial apical restriction of Baz and Armadillo (Arm= ßcat) during cellularization was disrupted (Fig S3I,J vs K,L, green vs yellow arrows), but this was largely restored at gastrulation onset (Fig S3M vs N), as is true in *cnoM/Z* null mutants. Thus, the RA domains are important for both Cno localization and function during initial apical-basal polarity establishment. This reinforced our only previous analysis of domain mutants, in which we examined cellularization after overexpressing CnoΔRA using the GAL4 system in a *cno* RNAi background (Bonello et al., 2018).

Cno plays a key role in ensuring robustness of AJs challenged by cell shape change/rearrangements. Cno is thus important for effective mesoderm apical constriction, the first gastrulation event. Mesoderm invagination of *cnoΔRA* mutants often did not go to completion (Fig 2A vs B; with 20/41 expected to be *M/Z* mutants, 27/41 embryos had incomplete closure at stages 7-8; Fig 2C, red arrow). This phenotype was similar to, though less severe than, that of *cnoM/Z* nulls (Fig 2D; Sawyer et al., 2009).

**Fig 2.**
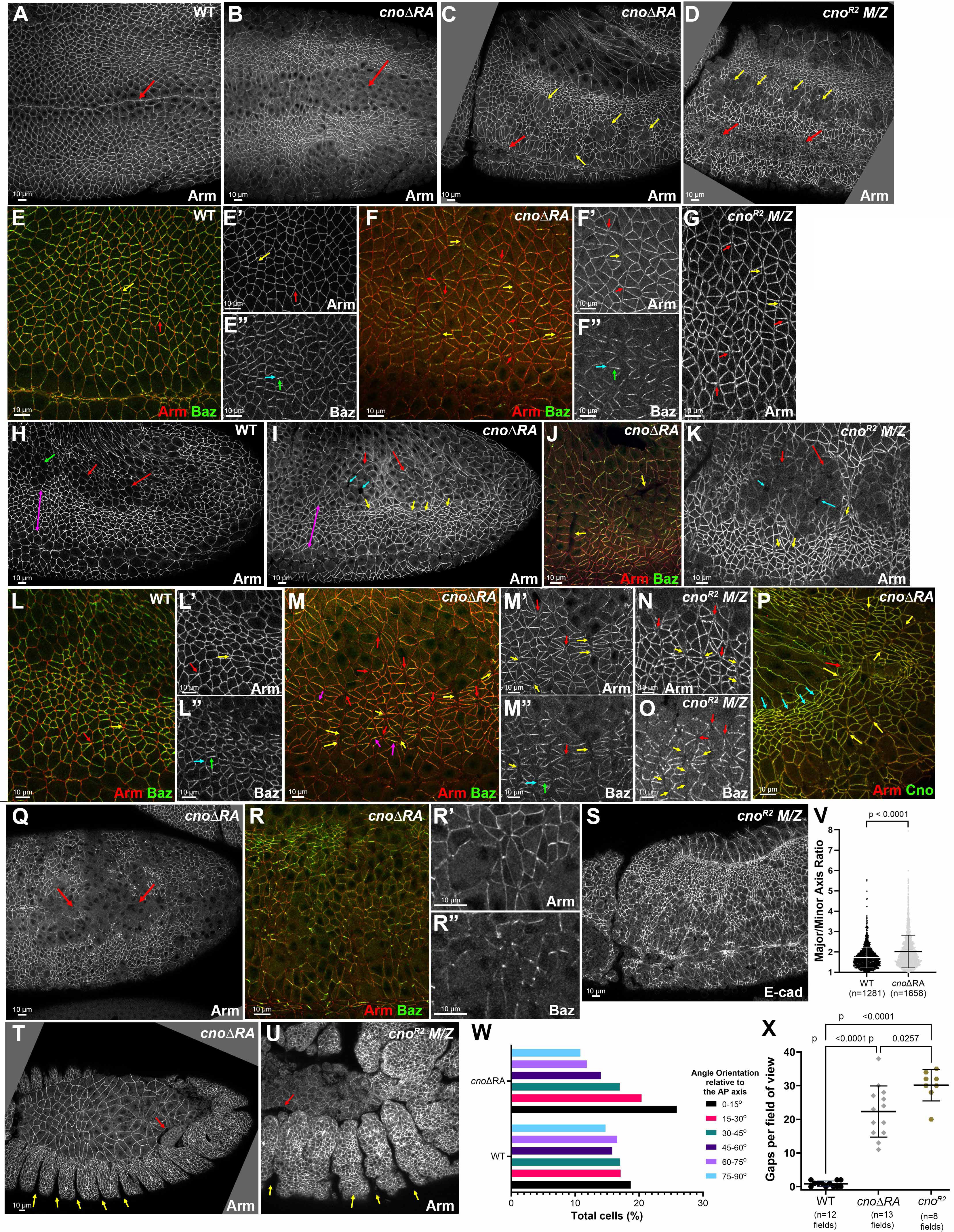
Deleting the RA domains substantially reduces Cno function in morphogenesis. Embryos, genotype and antigens indicated. In all Figs embryos are anterior left and, unless noted, dorsal up. A,B. Stage 8 ventral view. Dorsal closure is delayed in *cnoΔRA* (A vs B, arrows). C,D. Stage 9. *cnoΔRA* ventral open phenotype (C, red arrow) is similar to but less severe than *cno^R2^* (D, red arrows). In both more cells remain rounded up after division (yellow arrows). E-G. Stage 7. E. WT. Arm is continuous around cells (E, yellow arrow) and extends to TCJs (E, red arrow). F. *cnoΔRA.* Gaps at AP borders (F, yellow arrows) and TCJs (F, red arrows). Accentuated Baz planar polarity (E” vs F”, blue versus green arrows). G. Similar gaps in *cno^R2^*. H-O. Stage 8. H. WT. I,J.*cnoΔRA*. Neurectoderm (magenta bracket). Note stacks of elongated cells (I,J yellow arrows). Gaps (blue arrows) between mitotic cells (red arrows). Cells folded inward (J, arrows). K. Similar defects in *cno^R2^*. L-O. Closeups, stage 8. L. WT. Arm is continuous at shrinking AP borders (yellow arrow) and TCJs (red arrow). M. *cnoΔRA*. Large gaps along many AP borders (yellow arrows) or at TCJs (red arrows). Baz is hyper-planar polarized (L” vs M”). K, N,O. *cno^R2^* has similar defects. P. Stage 9. *cnoΔRA*. Cells remain elongated and aligned in stacks (blue arrows). Gaps remain (yellow arrows), especially where tissue is most curved (red arrow). Q-R. Stage 10. Epithelial integrity disruption at the ventral midline (Q). Baz lost from AJs more rapidly than Arm (R). S. *cno^R2^* has similar defects. T,U. Stage 13. *cnoΔRA* (T). *cno^R2^* (U) Segmental grooves remain (yellow arrows). Epidermis separates from amnioserosa (red arrows). V,W,X. Cell shape, orientation and Gap quantification: mean±SD.

Germband elongation further illustrates Cno’s role. During stages 7-8 anterior-posterior (AP) borders shrink by myosin-based contractility to form T1 and rosette cell arrangements, with new dorsal-ventral (DV) border extension completing cell intercalation (Guillot and Lecuit, 2013; Pare and Zallen, 2020). During stage 8 AJs are further challenged by the AJ remodeling required as cells round up for mitosis or invaginate as neural stem cells. In Cno’s absence these cell rearrangements are seriously disrupted (Sawyer et al., 2011).

*cnoΔRA* defects strongly resembled those of *cnoM/Z* null mutants. While Arm and Pyd localized to AJs in *cnoΔRA* mutants (Fig 2A vs B; H vs I; Fig S1C), defects in epithelial integrity emerged as germband extension began. At stage 7 small apical gaps appeared between cells at TCJs (Fig 2E vs F, red arrows) and AP cell interfaces (Fig 2E vs F, yellow arrows), similar to defects in *cnoM/Z* null mutants (Fig 2G). During WT stage 8, cells round up to divide in programmed domains—most prominent is mitotic domain 11 (Fig 2H, red arrows). More ventral cells (Fig 2H, magenta arrow) yet to divide rearrange to extend the germband. AJ proteins localize to both AP and DV borders (Fig 2L, yellow arrow) and extend to rosette centers (Fig 2L, red arrow). In *cnoΔRA*, cell rearrangements and shapes were altered—stacks of cells elongated along the AP axis were common (Fig 2I, yellow arrows). Quantification confirmed cell elongation and preferential cell alignment along the AP axis (Fig 2V,W). Looking more closely, gaps appeared at the center of most rosettes in *cnoΔRA* mutants (Fig 2M, red arrows) and cells separated along AP boundaries of aligned cells (Fig 2M, yellow arrows), both places where force on AJs is highest. Gaps also appeared between dividing cells (Fig 2I, cyan arrow). At times, cells folded inward at AP borders or within mitotic domains (Fig 2J, arrows). *cnoM/Z* null mutants were qualitatively similar (Fig 2K; Sawyer et al., 2011), with gaps at AP borders and rosette centers (Fig 2N, yellow and red arrows). Quantification verified the strong defects at TCJs and AP borders in *cnoΔRA* (Fig 2X)— there were 0.8 gaps per 133 x 133 µm field of cells in WT (n=12 embryos) vs. 22.3 gaps in *cnoΔRA* (n=13 embryos). *cnoM/Z* nulls were somewhat more severe, with 30.1 gaps (Fig 2X). Thus the RA domains are critical for Cno’s role in reinforcing AJs under elevated tension, and their loss strongly reduces Cno function.

In Cno’s absence, overall epithelial integrity is largely maintained, with cells along the ventral midline most sensitive to Cno loss (Sawyer et al., 2009; Manning et al., 2019). Once again, loss of the RA domains strongly reduced Cno function. In *cnoΔ*RA mutants, rows of aligned cells remained prominent during stage 9, as did gaps between them (Fig 2P, blue and yellow arrows, respectively)—this often was accompanied by tissue folding, particularly near the posterior end of the egg (Fig 2P, red arrow; 11/17 embryos). Cells rounded up for mitosis had delays in or failure to resume columnar architecture (Fig 2C, yellow arrows; 20/41 embryos, consistent with the 50% M/Z mutants), as in *cnoM/Z* null mutants (Fig 2D, yellow arrows; (Sawyer et al., 2009) or after *cno* RNAi (Manning et al., 2019). Defects were most common along the ventral midline (Fig 2Q; 6/20 presumptive *cnoΔRA M/Z* mutants). Baz was often lost or fragmented before Arm/AJs were lost (Fig 2R-R”), another phenotype of *cnoM/Z* null mutants. The most severe epithelial disruption phenotypes were less common in *cnoΔ*RA than in *cnoM/Z* nulls (Fig 2S), but many *cnoΔ*RA embryos had dorsal closure defects, including separation of the amnioserosa and epidermis (Fig 2T, red arrow) and retention of very deep segmental grooves (Fig 2T, yellow arrows). These were similar to though less severe than those of *cnoM/Z* null mutants (Fig 2U). Thus the RA domains play critical roles in every morphogenetic event in which Cno is implicated. This strong loss of function suggested *cnoΔ*RA provided a reasonable background in which to introduce our rescue constructs.

### The RA domains are required for Cno recruitment to AJs under tension

In both fly embryos and cultured mammalian cells Cno/Afadin accumulate at AJs under elevated tension.

During germband extension, these include AP borders and TCJs. CnoΔRA recruitment to these sites is dramatically altered. While WT Cno is mildly enriched on AP borders (Manning et al., 2019; Fig 1H), CnoΔRA accumulation on AP borders is strongly reduced, leading to planar polarization to DV borders (Fig 1I; quantified in Fig 1J). This suggests the RA domains are required for recruitment to AP borders, perhaps in response to elevated tension. Loss of the RA domains also nearly eliminated Cno TCJ enrichment during stage 7 (Fig 3A vs B; quantified in F), also sites of elevated tension (Yu and Zallen, 2020). Thus the RA domains are important for Cno recruitment to AJs under elevated tension.

**Fig 3.**
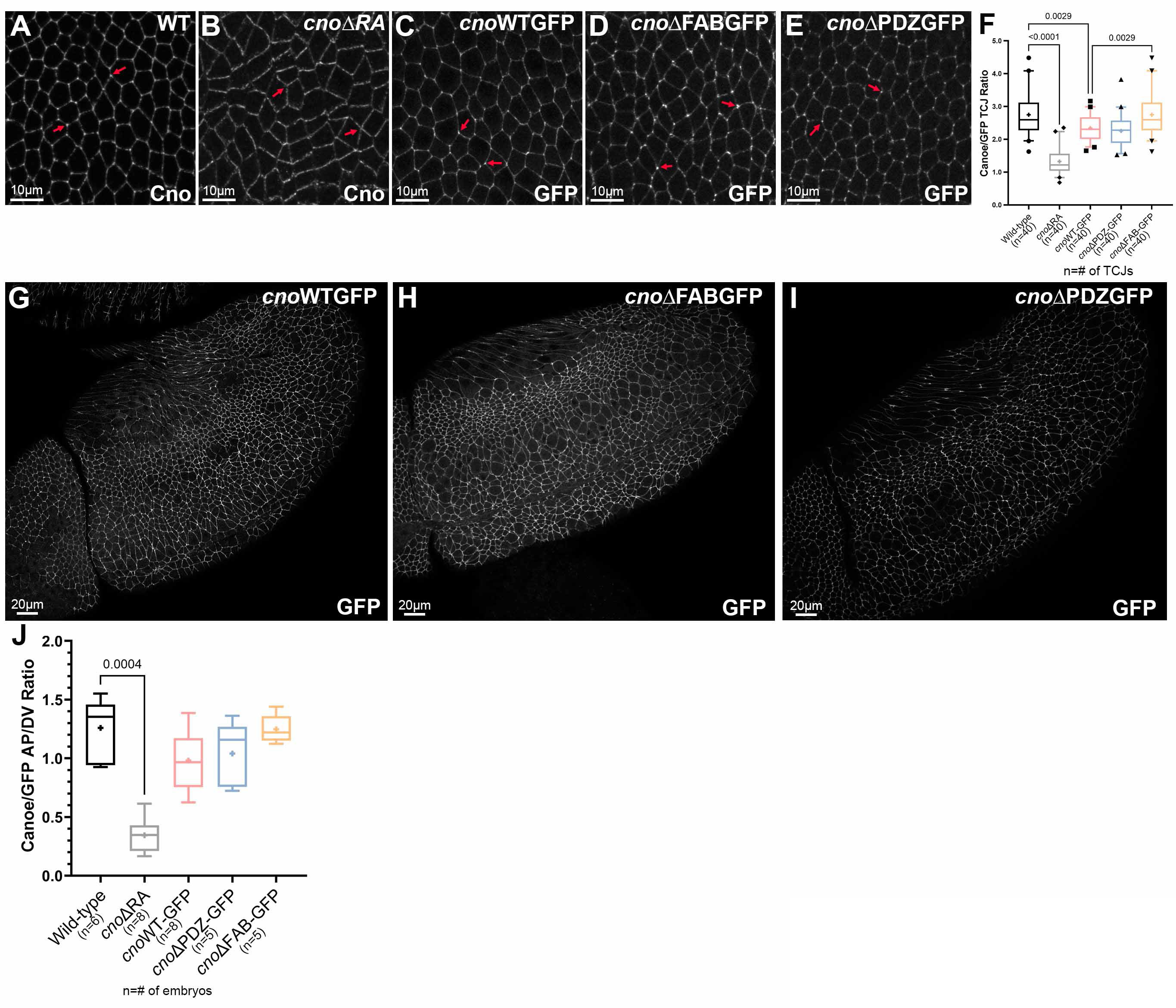
The RA, PDZ, and FAB domains are dispensable for Cno localization to AJs, and only the RA domain is important for TCJ enrichment. A-E. Stage 7. WT Cno, CnoWT, CnoΔFAB, and CnoΔPDZ all are enriched at TCJs (arrows). CnoΔRA TCJ enrichment is substantially reduced. F. Quantification. TCJ enrichment. G-I. Stage 9. CnoΔFAB and CnoΔPDZ AJ localization is unchanged. J. Quantification. Cno planar polarity.

Germband extension requires reciprocal planar polarization of AJ proteins and Baz on DV borders (Fig 2E, Baz in green) and actin, myosin and Cno on AP borders (Bertet et al., 2004; Zallen and Wieschaus, 2004). Cno restrains Baz and AJ planar polarization, preventing their loss from AP borders (Fig 2O; Sawyer et al., 2011). In *cnoΔ*RA, Baz planar polarity was strongly accentuated, as Baz was lost from AP borders (Fig 2E” vs F”, cyan arrow; Fig 2L” vs M”; quantified in Fig S1D) and sometimes concentrated at the center of DV borders (Fig 2M, magenta arrow). Both phenotypes occur in *cnoM/Z* null mutants (Sawyer et al., 2011). AJ planar polarization was also enhanced (Arm; Fig S1D), as in *cnoM/Z* null (Sawyer et al., 2011), as was planar polarity of the ZO-1 homolog Pyd (Fig S1D). Thus, the RA domains are important for Cno to regulate AJ/Baz planar polarity.

### Defining Cno PDZ domain structure

Two other Cno protein domains (Fig 4A) have defined binding partners: the PDZ and FAB. They could provide a simple linear link between AJs and actin. Scientists solved the structure of Afadin’s PDZ domain bound to C-terminal peptides of two known ligands: Nectin-3 (Fujiwara et al., 2015) and Bcr (Chen et al., 2007). The C-terminal tail of fly Ed, a nectin relative, can bind Cno’s PDZ (Wei et al., 2005). To gain molecular insight into Cno PDZ function, we crystallized the complex and determined its structure, fusing Ed’s C-terminal tail to the PDZ domain, enabling Ed-PDZ binding (Fujiwara et al., 2015). The structure was refined to 2.1 Å, representing the first Cno PDZ structure determined (Table 1 has data collection and refinement statistics). The structure adopts canonical PDZ architecture with a 5-stranded β-sheet flanked by two α-helices collectively forming a hydrophobic binding pocket containing the Ed peptide (Fig 4B,C; the flexible linker is not shown).

**Fig 4.**
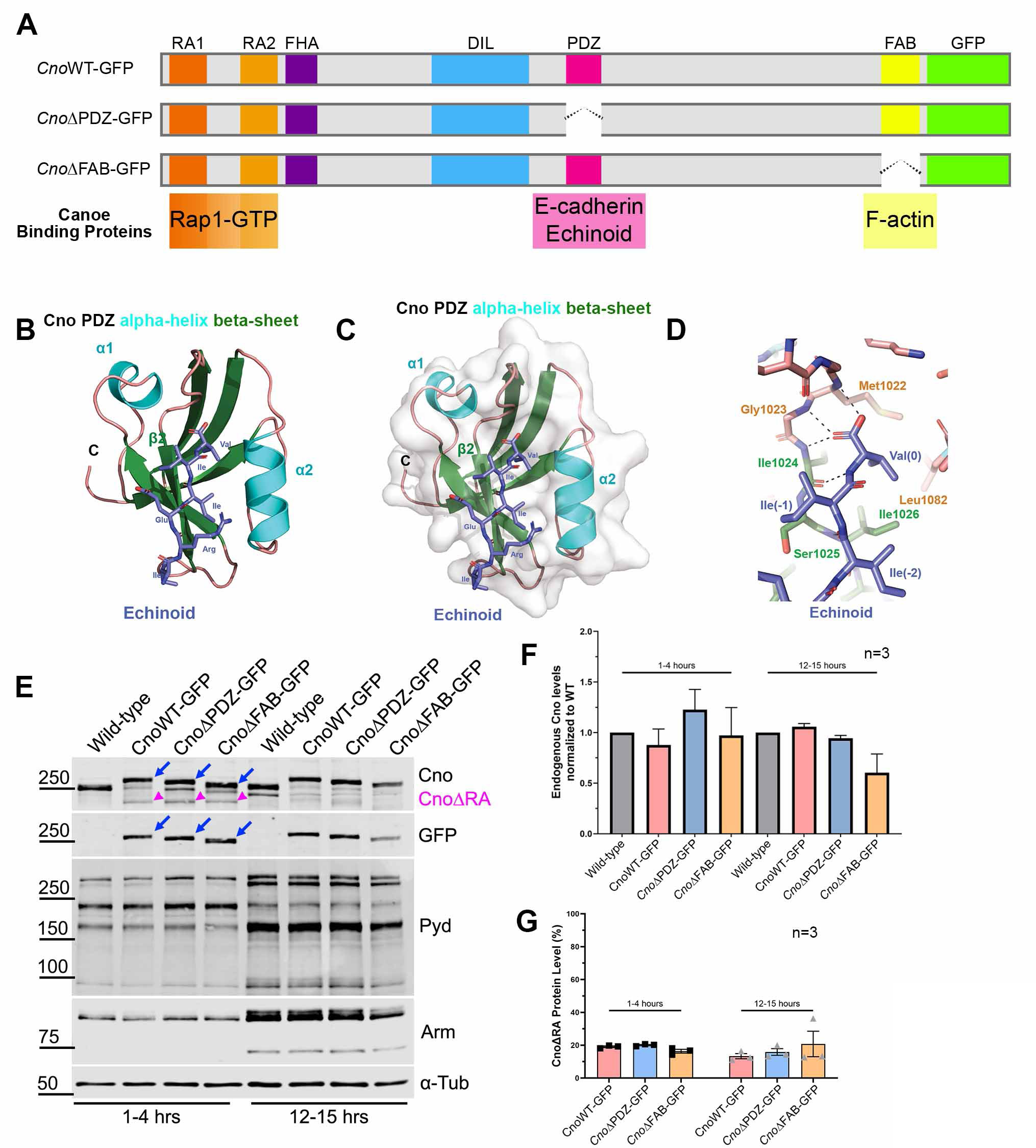
Defining Cno’s PDZ domain structure and generating mutants deleting it or the FAB. A. Cno protein domains, CnoΔPDZ and CnoΔFAB. B. Ribbon diagram, Cno PDZ (Green=β-strands, Cyan=α-helices; Teal=loops) bound to Ed’s C-terminal peptide (purple; IREIIV-COOH). C. Surface structure. D. Zoom view. Cno PDZ:Ed binding site. Key hydrophobic residues in binding pocket form van der Waals contacts with Ed and backbone determinants in the binding groove form hydrogen bonds (black dashed lines) with Ed’s terminal valine. E. Mutants are expressed at WT levels. Immunoblot, embryonic extracts. *α*-tubulin=loading control. Blue arrows=full length mutant proteins. Magenta arrowheads=truncated CnoΔRA. F. Protein levels relative to WT. G. Levels of residual CnoΔRA.

**Table 1.** Data processing and refinement statistics.

| **Crystal** | **Canoe PDZ-Echinoid** |
| --- | --- |
| **Data Collection** |  |
| Wavelength (Å) | 1.000 |
| Space group | P6 |
| Cell dimensions: a, b, c (Å) | 114.6, 114.6, 31.6 |
| Resolution (Å) | 50.00-2.10 (2.18-2.10) |
| # Reflections: Measured /<br>Unique | 154,188 (13,954) / 15,368<br>(1360) |
| Completeness (%) | 98.6 (99.3) |
| Mean redundancy | 11.0 (11.3) |
| $\langle I/\sigma I \rangle$ | 42.2 (6.9) |
| R<sub>sym</sub> | 0.074 (0.497) |
| CC1/2 | (0.966) |
| CC* | (0.991) |
| **Refinement** |  |
| Resolution (Å) | 37.5-2.10 (2.18-2.10) |
| R/ R<sub>free</sub> (%) | 19.2 (23.5) / 22.6 (26.8) |
| # Reflections, R/R<sub>free</sub> | 12,465 (1204) / 1,388 (136) |
| Total atoms: Protein / Water | 1405 / 112 |
| Stereochemical ideality<br>(rmsd): Bonds (Å) / Angles<br>(°) | 0.003 / 0.724 |
| Ramachandran Analysis:<br>Favored / Allowed (%) | 97.8 / 2.2 |
| **PDB accession code** | 7MFW |

Values in parentheses indicate statistics for the highest-resolution shell.

Within this pocket, Ed’s terminal valine residue’s carboxyl group forms hydrogen bonds with the backbone of Cno Met1022, Gly1023, and Ile1024. The side chains of this valine (position 0) and the isoleucine at position -2 form van der Waals contacts with hydrophobic residues lining the binding pocket (Fig 4D; Fig S4B,C). Ed binding buries 307 Å^2^ of PDZ solvent accessible surface area, and 393 Å^2^ of Ed C-terminal tail solvent accessible surface. Binding pocket surface residues are highly conserved (Fig S4A asterisks), with conservation extending to regions flanking the α-1 helix. Overall, Cno’s PDZ structure is highly similar to apo and peptide-bound structures of mammalian Afadin PDZs (Chen et al., 2007; Fujiwara et al., 2015; Joshi et al., 2006; Zhou et al., 2005). When peptide-bound Cno/Afadin PDZ structures are aligned, variation in PDZ sequence and plasticity become apparent (Fig S4D). While the ultimate residue in each target peptide is valine, residues at position -2 vary (Ed: isoleucine, Bcr: threonine, Nectin-3: tryptophan). Interestingly, as the side chain of residue -2 increases in size, there is a corresponding shift of the α2-helix outwards, effectively opening the binding pocket to accommodate the larger residue. The Cno PDZ-Ed structure highlights a key biological interaction and primed us to investigate effects of deleting the PDZ on Cno function.

### The PDZ and FAB are not required for viability, but support viability and morphogenesis when protein levels are reduced

We next directly tested the hypothesis that the PDZ and FAB play key roles in AJs:cytoskeletal connections. To do so, we used the site-specific recombination site to re-introduce CnoΔPDZ and CnoΔFAB, which cleanly deleted each domain, in parallel with a WT Cno construct (CnoWT; Fig 4A). The crystal structures guided PDZ deletion (94 amino acids deleted), and for the FAB sequence conservation guided deletion of the 115 most conserved amino acids (Fig S4E). All restore missing intron sequences, the 5’ UTR and start codon, WT or mutated coding sequence, a C-terminal GFP-tag, the 3’ UTR and polyA signal. We first used immunoblotting to verify that embryos homozygous for these constructs expressed a GFP-tagged protein of the predicted sizes at WT levels (Fig 4E, blue arrows; quantified in Fig 4F). Importantly, inserting these constructs into the locus restored splicing of exon 1 to re-introduced exon 2, substantially reducing levels of the truncated CnoΔRA protein produced by mis-splicing to 13-21% of total protein (Fig 4E, magenta arrowheads; quantified in Fig 4G). This further reduced concerns about residual function of CnoΔRA, which serves as the background.

CnoWT, CnoΔPDZ, and CnoΔFAB enrichment at AJs resembles WT Cno (Fig 3G-I). We thus tested CnoΔPDZ and CnoΔFAB function, with CnoWT as our control.

*cno* null mutants are zygotically lethal, and reintroducing CnoWT restored zygotic and M/Z viability and fertility (Fig 5A), verifying our “knock-in” rescue strategy. We expected neither CnoΔPDZ nor CnoΔFAB would rescue viability. However, to our surprise, homozygous zygotic *cnoΔPDZ* and *cnoΔFAB* mutants survived to adulthood (Fig 5B,C), and even more surprising, we were able to generate viable, fertile stocks of each mutant. Thus neither the PDZ nor the FAB is required for viability, contrary to the first hypothesis. This result was sufficiently surprising that we recreated the *cnoΔFAB* mutant via CRISPR directly in the endogenous locus, without the *cnoΔRA* deletion. These new *cnoΔFAB* mutants were also viable and fertile, and did not produce CnoΔRA protein (Fig S5). We next assessed embryonic viability of M/Z *cnoWT, cnoΔPDZ* and *cnoΔFAB* mutants, as a stock can be maintained despite measurable embryonic lethality. Strikingly, M/Z

**Fig 5.**
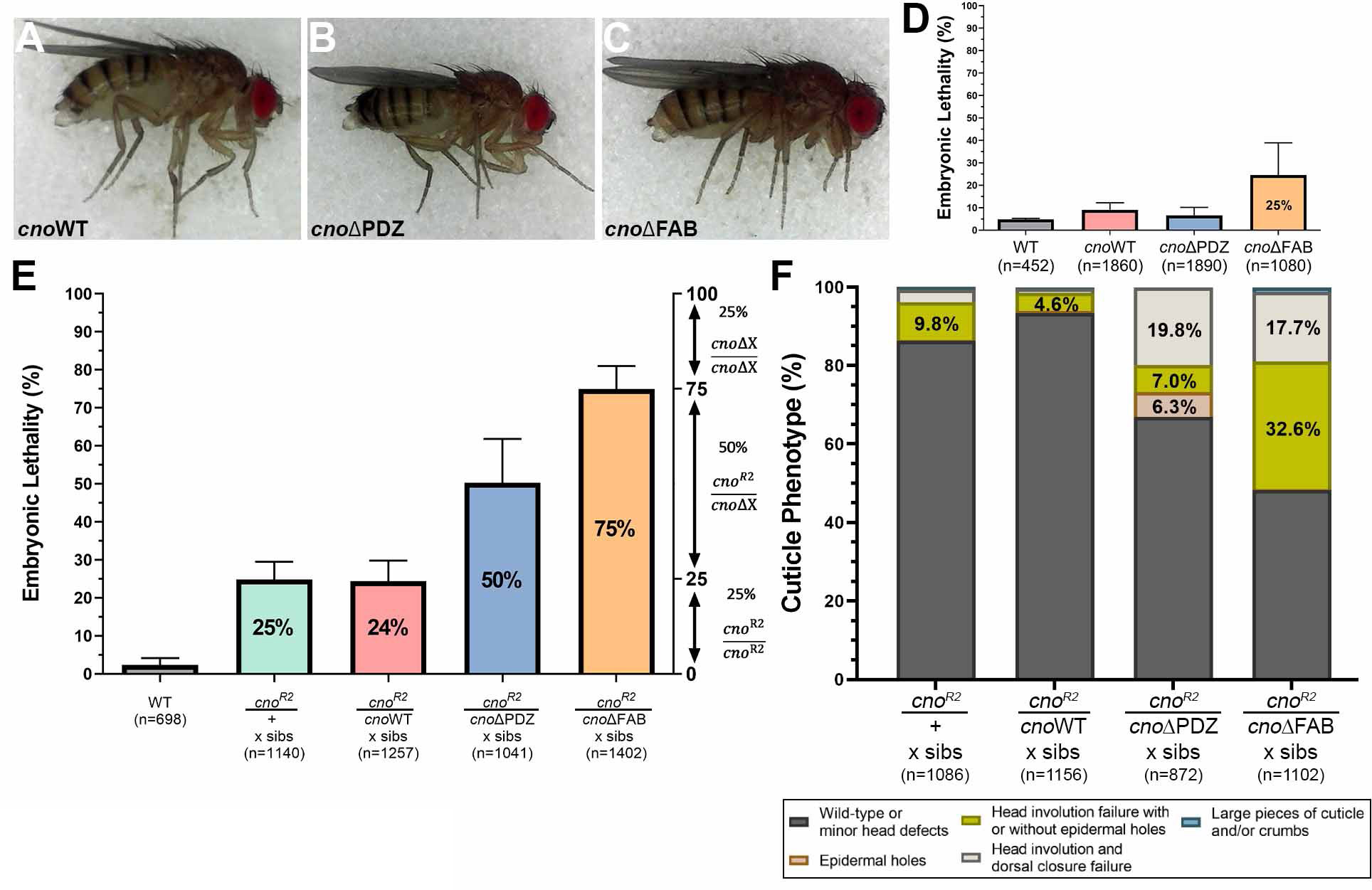
Neither the PDZ nor the FAB are required for viability, but sensitized assays reveal roles in morphogenesis. A-C. Viable homozygous adults. D. Embryonic lethality, M/Z mutants. E. Sensitized assay revealed reduced function of CnoΔPDZ and CnoΔFAB. Embryonic lethality, progeny of crosses of WT or *cno* mutants heterozygous with *cno^R2^*. *cnoWT* behaves like a WT allele while *cnoΔPDZ,* or *cnoΔFAB* provide less function. F. Cuticles reveal reduced function in morphogenesis.

*cnoWT* and *cnoΔPDZ* mutants had normal embryonic viability, in the range of those seen in WT controls (Fig 5D). *cnoΔFAB* did not fully rescue viability, with 25% embryonic lethality (Fig 5D). In cuticles of the subset of *cnoΔFAB* embryos that died, head involution failed in 42% (n=124), consistent with the fact that head involution is the event most sensitive to reduced *cno* function (Sawyer et al., 2009). Thus, while the FAB is not essential, it is required for full protein function.

To increase assay stringency, we used genetic crosses to reduce mutant protein levels, by making mutants heterozygous with the null allele, *cno^R2^.* This proved quite revealing. *cno^R2^* is zygotically embryonic lethal when homozygous (Sawyer et al., 2009), and thus when one crosses *cno^R2^/ +* females and males embryonic lethality should be 25%, with only *cno^R2^/ cno^R2^* progeny dying (we measured 24.8%; Fig 5E; Fig S2). *cno^R2^/ +* (50% of progeny) and *+/+* progeny (25% of progeny) should survive. If our *cno* mutant alleles were fully functional, they should behave similarly to a WT chromosome. When we crossed *cno^R2^/cnoWT* females and males, that is what we saw: 24.4% lethality (Fig 5E). In contrast, when we crossed *cno^R2^/cnoΔPDZ* females and males there was 50.3% lethality, suggesting some *cno^R2^/cnoΔPDZ* progeny also die as embryos (Fig 5E). This effect was even more striking in the *cno^R2^/cnoΔFAB* cross, with 74.9% lethality, consistent with all *cno^R2^/cnoΔFAB* progeny dying as embryos and only the 25% *cnoΔFAB* homozygous embryos surviving (Fig 5E). We saw very similar trends when we crossed *cno^R2^/mutant* females to males homozygous for our site-directed mutants (Fig S2)— *cnoWT* fully rescued embryonic viability (7% lethal, similar to WT), *cnoΔPDZ* was less functional (23.6% lethality) and *cnoΔFAB* least functional (62.0% lethality).

Larval cuticle analysis reinforced these data. Most progeny from the *cno^R2^/cnoWT* self cross were either WT or had defects in head involution (Fig 5F), similar to *cno^R2^/ cno^R2^* zygotic mutants (Sawyer et al., 2009). Progeny from the *cno^R2^/cnoΔPDZ* self cross had more severe phenotypes with 19.8% exhibiting defects in both dorsal closure and head involution (Fig 5F). Progeny from the *cno^R2^/cnoΔFAB* self cross had even more severe cuticle phenotypes (Fig 5F). Thus, while the PDZ and the FAB are not absolutely required for viability, they become important for rescuing viability and morphogenesis when mutant protein levels are reduced, with the FAB making a more substantial contribution to protein function than the PDZ.

### The FAB domain is important to maintain full junctional integrity at TCJs

Analysis of *cnoM/Z* null mutants (Sawyer et al., 2009; 2011) and *cnoΔRA* (above) revealed important roles for Cno in embryonic morphogenesis. However, other mutants affecting morphogenesis have more subtle defects. Especially notable is the TCJ protein Sdk; mutants are viable and fertile but have subtle defects in cell shape and in AJ integrity at TCJs, where Sdk localizes (Finegan et al., 2019). We wondered whether *cnoΔFAB* or *cnoΔPDZ* had similar defects compatible with viability. To explore this, we visualized AJ proteins in *cnoΔFAB* and *cnoΔPDZ*.

We began with *cnoΔFAB,* as embryonic lethality and cuticle data suggested the FAB plays a more important role in morphogenesis than the PDZ. Both CnoWT (Fig 6D-F) and CnoΔFAB properly localized and functioned during cellularization. Like WT Cno, CnoΔFAB protein accumulated in nascent AJs and not basal junctions (Fig 6A,C vs J,L yellow versus red arrows; quantified in Fig 6P vs R) and was enriched at TCJ cables extending below nascent AJs (Fig 6B’ vs K’, yellow arrows). *cnoΔFAB* mutants had normal Arm localization (Fig 6A,C vs J,L yellow versus red arrows; quantified in Fig 6M vs O). Mesoderm invagination requires Cno (Sawyer et al., 2009), and the first phenotype in *cnoΔFAB* mutants occurred during this process. In *cnoΔFAB* 14/21 stages 7-9 embryos had mesoderm cells still present at the embryo surface (Fig 7A vs B). However, these defects were less severe than those of *cnoΔRA* or *cnoM/Z* null mutants, resolving as development proceeded.

**Fig 6.**
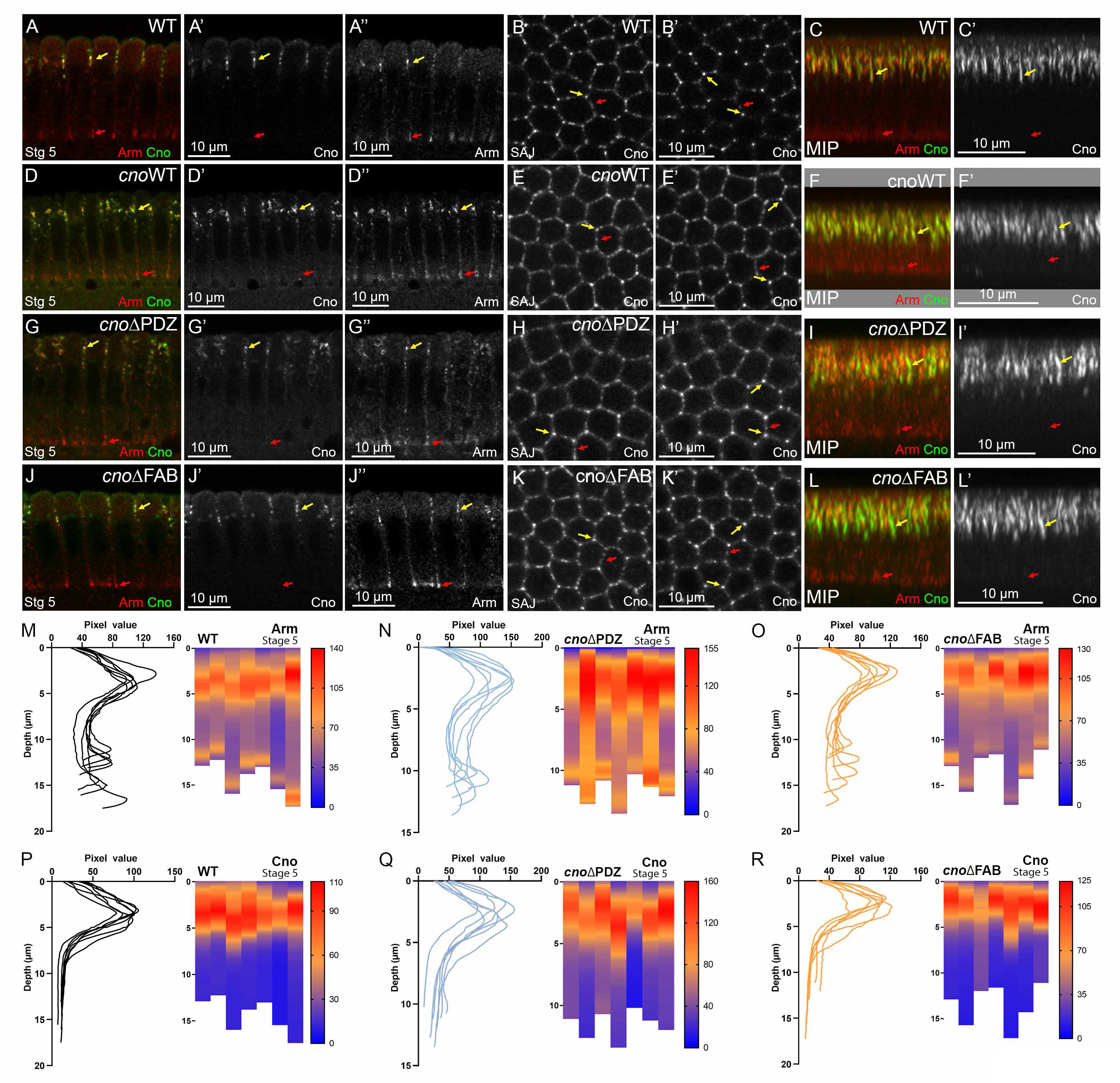
The PDZ and FAB are not required for Cno localization/function during cellularization. A-O. Stage 5. A,D,G,J. Cross-sections. B,E,H,K. En face. Spot AJs (SAJ). B’,E’,H’,K’. 33.33% below SAJs. C,F,I,L. Maximum intensity projection (MIP). M-R. Line traces and heat map quantification of Arm (M-O) or Cno (P-R) mean intensity along lateral membranes. Each column is an embryo (n=7). A-C. WT. Cno and Arm are enriched apically (A-C, yellow arrows). Arm is also at basal junctions (red arrows). Cno localizes to bicellular and TCJs at the level of SAJs (B) and is enriched at TCJs deeper into embryos (B’, C). D-L. CnoWT (D-F), CnoΔPDZ (G-I), and CnoΔFAB (J-L) localize like WT Cno. All also have normal Arm enrichment in apical SAJs and basal junctions.

**Fig 7.**
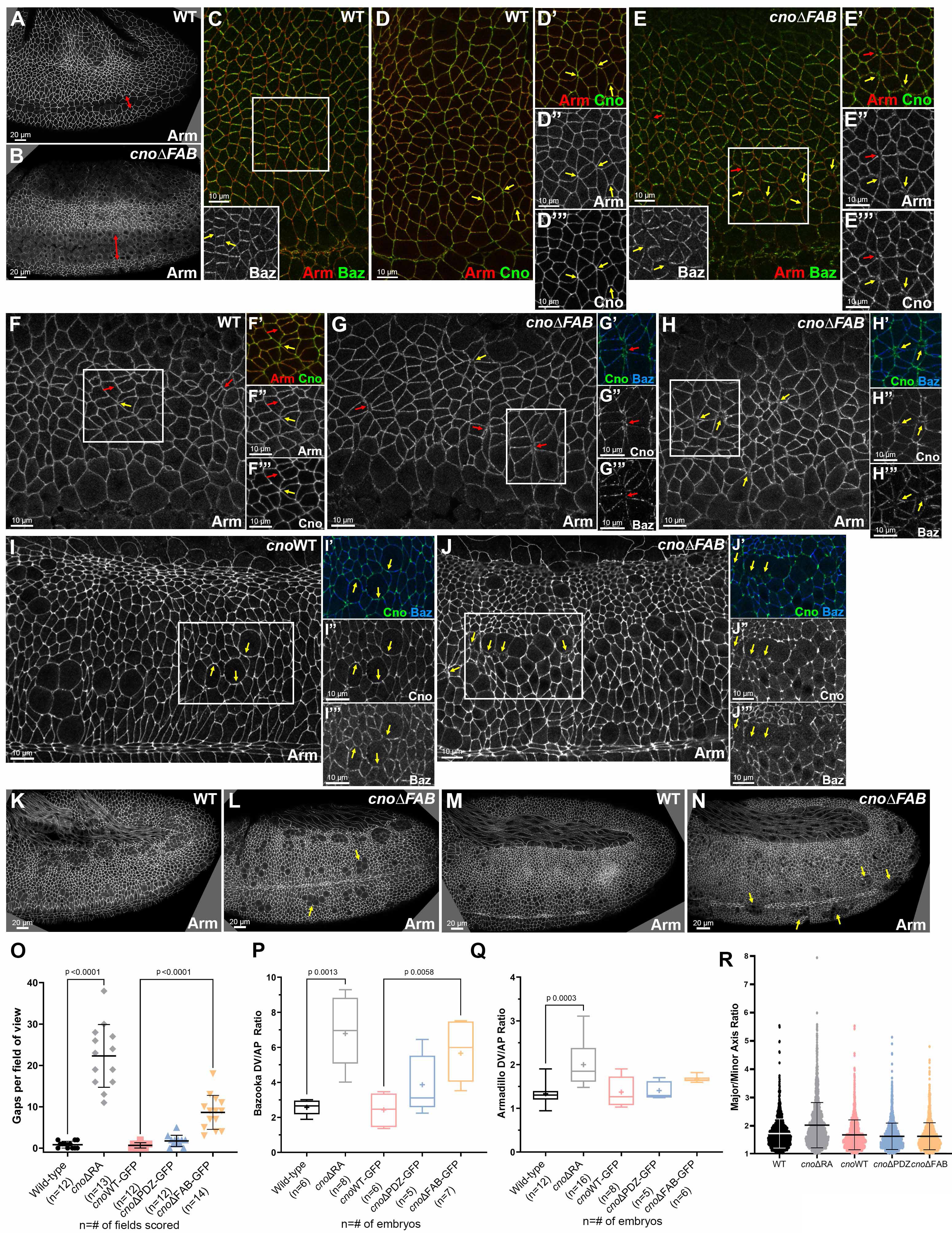
Deleting the FAB leads to defects in mesoderm invagination and at AJs under tension. A,B. Stage 8. *cnoΔFAB* has impaired mesoderm invagination (arrows). C-E. Stage 7. WT Arm and Cno extend to TCJs (D, yellow arrows). Baz is planar polarized (C inset, arrows). E. *cnoΔFAB.* Small gaps (E-E”’, red arrows) or fragmented TCJs (yellow arrows). Accentuated Baz planar polarity with loss from AP borders (E inset, yellow arrows). F-H. Stage 8. WT Arm and Cno are continuous at shrinking AP borders (F, red arrows) and TCJs (F, yellow arrow). *cnoΔFAB*. Gaps at TCJs (G,H yellow arrows) and along AP borders of aligned cells (F vs G red arrows). Baz is lost there (G, inset). I-J. Stage 9. *cnoΔFAB*. Modest defects in AJs remain near mitotic cells (I vs J, yellow arrows). K-L. Stage 10. M-N. Stage 11. *cnoΔFAB.* Cells along the ventral midline delayed in resuming columnar architecture (L, arrows) or with reduced epithelial integrity (N, arrows). O-R. Quantification, Gaps., Baz and Arm planar polarity, cell elongation.

During gastrulation CnoΔFAB protein localized like WT Cno and CnoWT (Fig 3G vs H), localizing to AJs, enriched at TCJs (Fig 3A,C vs D, quantified in F), and without the reversed planar polarization of CnoΔRA (Fig 3J), suggesting it still is recruited to AJs under elevated tension. As germband extension began, *cnoΔFAB* mutants did not exhibit the dramatic early disruption of epithelial integrity of *cnoM/Z* null or *cnoΔRA* mutants. However, they clearly deviated from WT in AJ stability. During WT stage 7, as cells began T1/rosette rearrangements, Baz is planar polarized to AP boundaries (Fig 7C, inset), while Arm and Cno extend around the circumference with Cno enriched at TCJs (Fig 7D, arrows). *cnoΔFAB* defects appeared at this stage, with gaps (Fig 7E, red arrows) or Arm and Cno fragmentation/broadening at TCJs (Fig 7E, yellow arrows). Baz was lost from some AP borders (Fig 7E, inset), elevating planar polarization (Fig 7P), though not as dramatically as in *cnoΔRA*. There was subtle elevation of AJ planar polarity (Fig 7Q), but the cell elongation and alterations in Pyd planar polarity seen in *cnoΔRA* were absent (Fig 7R; S1C vs E-G). Defects in AJ integrity and Baz planar polarity continued into stage 8. In WT, Arm localized to AP and DV AJs (Fig 7F, red arrows), and extended to TCJ centers (Fig 7F yellow arrow). In contrast, in *cnoΔFAB*, cell separation along aligned AP boundaries (Fig 7G, red arrows) and TCJ gaps or AJ broadening continued (Fig 7G,H, yellow arrows). Once again, quantification reinforced this (Fig 7O)— there were 8.6 gaps per field in *cnoΔFAB* (n=14 embryos) vs. 0.83 gaps per field in wildtype (n=12 embryos). Importantly, CnoWT embryos did not have defects in this assay (0.67 gaps per field; n=12 embryos). Small gaps or disruption at TCJs persisted in stage 9, most often near cells rounded up for division (Fig 7I vs J, arrows). By stage 10, the ectoderm was largely intact in *cnoΔFAB*, but groups of cells were delayed in resuming columnar architecture (Fig 4L, arrows) or had lost epithelial architecture, generally near the ventral midline (Fig 7N, arrows; 10/25 embryos had one of these defects). Thus, Cno’s FAB is not required for Cno recruitment to AJs under tension, but it is important for reinforcing those AJs. However, most cells recover, suggesting redundant interactions mediate Cno action.

### The PDZ only plays modest roles in AJ stability and maintaining columnar cell architecture

We next compared *cnoΔPDZ* to *cnoΔRA* and *cnoΔFAB*. During cellularization, CnoΔPDZ properly localized and rescued function. CnoΔPDZ was restricted to nascent AJs (Fig 6A,C vs G,I yellow versus red arrows; quantified in Fig 6P vs Q), and enriched in TCJ cables (Fig 6B’ vs H’, yellow arrows). *cnoΔPDZ* mutants had normal Arm localization (Fig 6A,C vs G,I yellow versus red arrows; quantified in Fig 6M vs N). In contrast with the other mutants, *cnoΔPDZ* mutants had no significant defects in mesoderm invagination (Fig 8A vs B; 21/22 embryos were normal). Germband extension proceeded normally, without defects in epithelial integrity (Fig 8C vs D). We closely examined embryos at stages 7-8, observing occasional gaps or AJ broadening at TCJs (Fig 8E,F, arrows), but their frequency was significantly lower than in *cnoΔFAB* mutants (Fig 7O). AJ (Fig 7Q) and CnoΔPDZ protein (Fig 3J) planar polarity were unaltered, there was only subtly enhanced Baz planar polarity (Fig 8F””; quantified in Fig 7P), and cells were not elongated (Fig 7R). During stages 9-10 most embryos looked WT (Fig 8G)—there was a slight increase in the frequency of embryos with multiple cells rounded up to divide simultaneously (Fig 8H—42% of *CnoΔPDZ* embryos (n=36) vs 30% of WT embryos (n=23))— consistent with a subtle delay in regaining columnar architecture after division. Embryos from stage 11 onward looked WT (Fig 8I vs J). Thus, when expressed at WT levels the PDZ is largely dispensable for most Cno’s roles and plays only a subtle role in ensuring AJ stability under mechanical stress.

**Fig 8.**
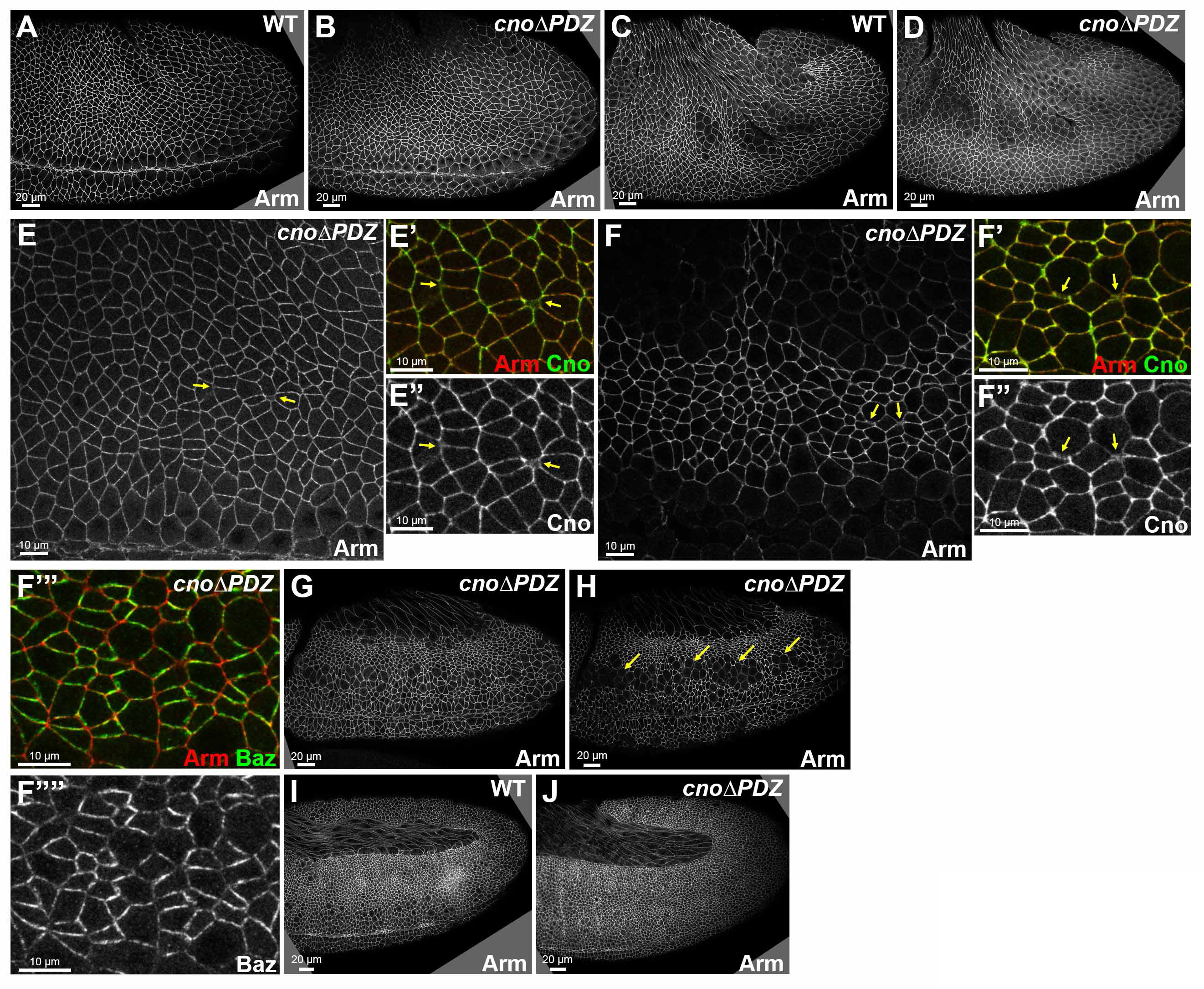
Deleting the PDZ has only modest effects on AJ integrity. A,B. Stage 7. *cnoΔPDZ*. No defects in mesoderm invagination. C,D. Stage 8. *cnoΔPDZ*. Epithelial integrity was largely normal. E. Stage 7. F. Stage 8. Occasional gaps or broadening of TCJs (arrows). Near normal Baz planar polarity (F’”,F ””). G,H. Stage 9. *cnoΔPDZ*. Subtle increase in embryos with multiple cells simultaneously rounded up to divide (H, arrows; 42% vs 30% in WT). I,J. Stage 10. *cnoΔPDZ* appeared normal.

## Discussion

One key issue for our field is defining mechanisms cells use to connect AJs to the cytoskeleton. This connection must be dynamic and force responsive to accommodate the many cell rearrangements and shape changes of morphogenesis. We focused on Cno, a critical part of this linkage, to determine its mechanism of action. Previous analyses of Cno function used null alleles. This approach limits our understanding of how Cno works as a machine to integrate multiple inputs and mediate AJ:actomyosin linkage, as Cno is a complex multidomain protein. Here we provide surprising new insights into Cno’s mechanism of action by interrogating the function of individual Cno domains. These analyses provide evidence that AJ:actomyosin linkage involves complex multivalent interactions conferring robustness and ensuring tissue integrity.

### A hierarchy of protein function, multivalent AJ assembly, and robustness

Diagrams of AJ:cytoskeletal linkage often suggested simple linear pathways of connection and function: e.g., cadherins recruit catenins which directly bind actin, or, in polarity establishment, Cno recruits Baz which recruits AJs proteins. Work in the last decade altered this view significantly. First, AJs are massive multiprotein assemblies—even at initial assembly spot AJs contain >1000 Ecad and >400 Baz proteins (McGill et al., 2009). Second, many proteins at the AJ-cytoskeletal interface are multidomain proteins interacting with multiple partners. This suggests a different view—the AJ:cytoskeletal interface assembles from a large protein network via multivalent interactions, with proteins serving as nodes in this network (Fig 9A), and multidomain proteins mediating multiple linkages (Fig 9B).

**Fig 9.**
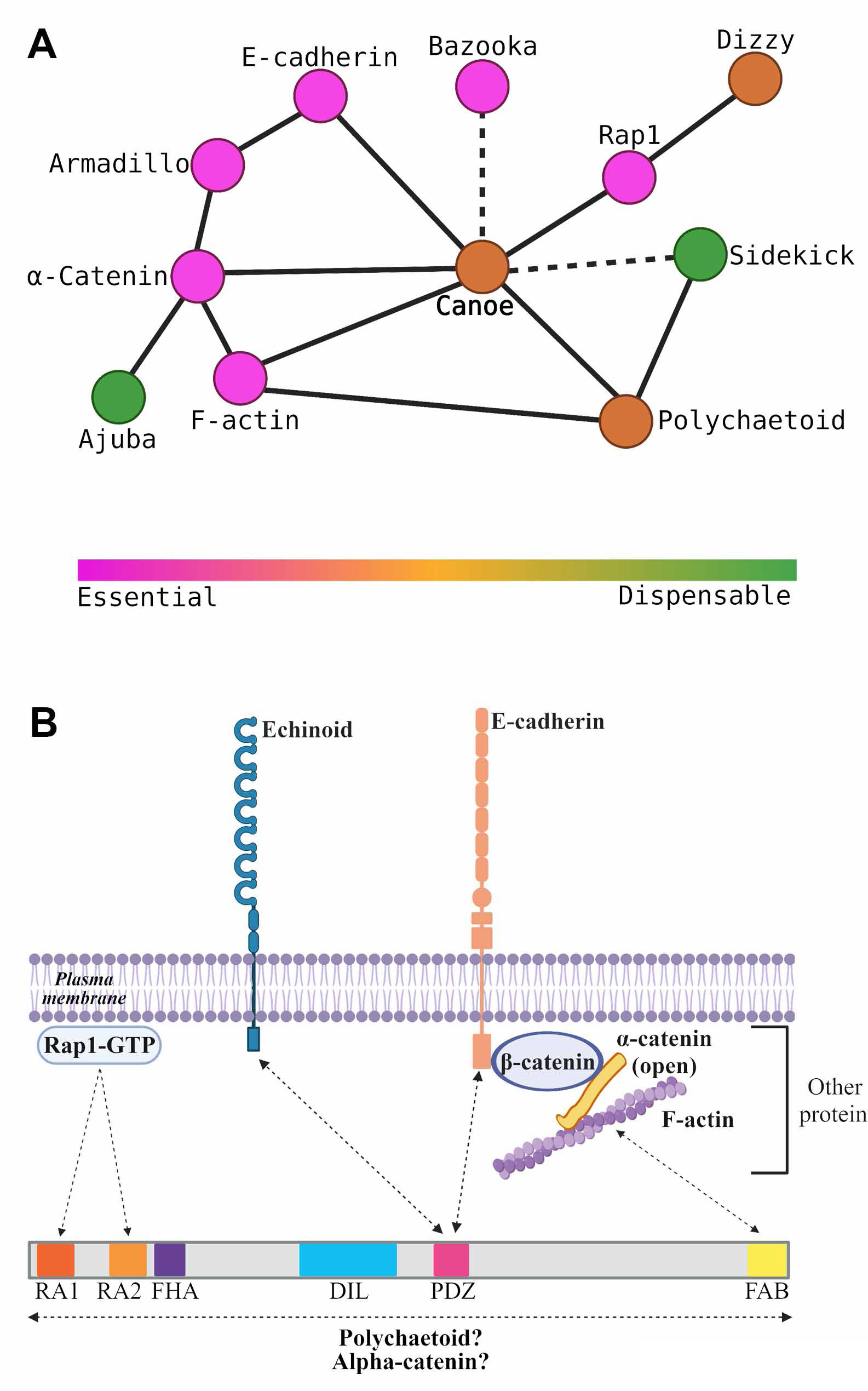
Two speculative models. A. Node model. Protein nodes range from essential (magenta) to dispensable (green). B. A network of protein assembled by multivalent interactions that provides tissue robustness.

In this AJ:cytoskeletal network, some interaction nodes are more central than others, revealing a hierarchy of proteins with different degrees of importance for epithelial integrity (Fig 9A). Ecad, the catenins, and Baz are essential for AJ formation and stability; without these, embryos fall apart at gastrulation (Cox et al., 1996; Müller and Wieschaus, 1996; Sarpal et al., 2012). Cno is not essential for cell adhesion, but in its absence there are defects in initial AJ polarization and in correct completion of many morphogenetic movements (Choi et al., 2013; Manning et al., 2019; Sawyer et al., 2011; Sawyer et al., 2009), resulting from disruption of cell shape change and AJ integrity in places where AJ forces are maximal. Pyd is zygotically viable, but 60% of M/Z mutants have defects in cell shape change during dorsal closure (Choi et al., 2011). Finally, Sdk is dispensable, but in its absence AJ integrity defects occur at TCJs (Finegan et al., 2019), where force exerted on AJs is high (Yu and Zallen, 2020). This hierarchy likely reflects the multivalent contacts within the AJ complex (Fig 9B), with more dispensable players located at more redundant nodes in the network. This view of differentially important nodes and interactions also may explain the differential effects of our Cno domain deletion mutants; we saw a functional hierarchy in which RA domain loss had the strongest effects, with FAB loss next and PDZ loss least severe. We saw similar effects on mammalian cell AJ integrity when rescuing Afadin shRNA knockdown with Afadin proteins lacking individual domains (Choi et al., 2016). Afadin lacking the RA domains had the most severe defects in tissue integrity, but no single domain deletion fully eliminated function. It will be exciting to explore the full extent of the AJ:cytoskeletal network, dissecting the importance of other nodes and connections.

### One key role for Cno is in reinforcing AJs under tension

This new network and node view of AJ:cytoskeletal connections helps illuminate Cno’s diverse roles. In Cno’s absence the cytoskeleton detached from AJs (Sawyer et al., 2011; Sawyer et al., 2009). During apical constriction, cell shape change began in Cno’s absence, with detachment occurring midway through constriction, suggesting Cno reinforcement is needed when AJs reach a critical tension threshold. Intriguingly, Cno’s homolog Afadin has a similar role in MDCK cells in which ZO-1 family protein knockdown elevated junctional tension. As in the embryo, the cytoskeleton lost its tight connection with AJs when Afadin was knocked down, with the weakest points at TCJs (Choi et al., 2016). Afadin is strongly recruited to AJs in response to elevated tension, suggesting it is recruited to reinforce AJ-cytoskeletal connections. In embryos, Cno is enriched at TCJs and AP borders (Bonello et al., 2018; Manning et al., 2019; Sawyer et al., 2011), both locations where tension is elevated (Fernandez-Gonzalez et al., 2009; Yu and Zallen, 2020). Inhibiting Rho-kinase leads to rapid Cno loss from TCJs, supporting the idea that Cno is recruited to TCJs by cytoskeletal tension (Yu and Zallen, 2020). Intriguingly, in ZO-knockdown MDCK cells, inhibiting myosin motor activity by blebbistatin also reduced Afadin AJ enrichment, while Rock inhibition fragmented junctional Afadin (Choi et al., 2016). Together these data suggest that a central role of Cno is to strengthen AJ:cytoskeletal connections under elevated tension, allowing them to respond to force exerted by actomyosin mediated contractility, and that Cno does so in a mechanosensitive way. Our data reveal new insights into this mechanism. Cno activation by Rap1 appears to play a critical role, as mechanosensitive Cno recruitment to TCJs and AP borders is lost in CnoΔRA, and CnoΔRA function in AJ reinforcement during mesoderm apical constriction and germband elongation is strongly impaired. The FAB and PDZ domains are less critical. Neither is essential for recruitment to TCJs. In this our data differ from that of Yu and Zallen, who suggested phosphorylation of a key tyrosine in the FAB by Abl kinase is important for Cno mechanosensitive recruitment (Yu and Zallen, 2020) —perhaps the difference reflects using GAL4-driven expression versus mutating *cno* at the endogenous locus. However, node connections made by the FAB are important to reinforce AJs under tension, leading to the gaps seen at TCJs and AP borders. In contrast, network connections made by the PDZ are dispensable, unless cells are sensitized by reducing Cno levels.

### Cno AJ recruitment involves multiple semi-redundant interactions

To function, Cno must localize to AJs in ways that are planar polarized and mechanically responsive. Our data build on previous work, revealing Cno localization is a complex response to multiple upstream regulatory cues and occurs by multivalent and often redundant interactions (Fig 9B). One important cue is the small GTPase Rap1. During cellularization, active Rap1 is necessary and sufficient to recruit Cno to AJs (Bonello et al., 2018). However, surprisingly, the RA domains that bind active Rap1 are not required for Cno cortical localization during cellularization, though they do mediate assembly into TCJ cables. Thus Rap1 regulates Cno localization by both RA-dependent and RA-independent means. Further, neither Rap1 nor the RA domains are required for Cno AJ localization after gastrulation onset (Bonello et al., 2018). However, our data reveal that the RA domains and presumably Rap1 input are required for Cno recruitment to AJs where tension is elevated. We still don’t know what it means mechanistically to say Rap1 “activates” Cno—e.g. does it open a closed conformation or mediate critical interactions? This is an important topic for future work. A second important cue for Cno localization is F-actin (Sawyer et al., 2009). The simplest explanation would be that this is mediated by Cno’s FAB, but, surprisingly, the FAB alone is not necessary for Cno AJ recruitment. CnoΔFAB is enriched at TCJs and AP borders, and thus the FAB is not required for mechanosensitive recruitment. These data suggest F-actin input into Cno localization occurs via multiple domains, likely via direct and indirect interactions.

Our data and work from the Zallen lab further emphasize the robust multivalent interactions mediating Cno recruitment. We were quite surprised that the PDZ, which binds the AJ transmembrane proteins Ecad and Ed, is not essential for AJ localization. Further, a Cno construct lacking the FHA, DIL and PDZ domains still localizes to AJs (Yu and Zallen, 2020), suggesting the RA domains, intrinsically disordered region and FAB can work together to mediate recruitment. Individual domains are less able to mediate localization— constructs encoding the RA domains or the intrinsically disordered region plus the FAB are only weakly recruited to AJs (Yu and Zallen, 2020), while a construct carrying the Cno FHA, DIL and PDZ domains is not recruited to AJs at all (Bonello et al., 2018). These data suggest recruitment is mediated by multivalent interaction, with no single interaction essential. Cno/Afadin can be recruited to AJs by active Rap1/RA, PDZ/Ecad, IDR/*α*cat or FAB/actin interactions, providing robustness to AJ assembly. Many AJ proteins like Pyd and Baz are also multidomain scaffolding proteins, and analysis of Baz similarly revealed deleting individual domains does not affect cortical localization (McKinley et al., 2012), suggesting this is a general feature of AJ scaffolding proteins.

## Acknowledgements

We are grateful to Andrew Spracklen for CRISPR/Cas9 advice, Nathaniel Wesley for protein production, Andrew Spracklen, Teresa Bonello, Lathiena Nervo, and other Peifer lab members for helpful advice and discussions, to Dan Bergstralh, Ulrich Tepass, Bob Duronio and Scott Williams for helpful discussions. We thank Noah Gurley and Melissa Greene for technical assistance, and Tony Perdue of the Biology Imaging Center for confocal imaging advice. This work was supported by NIH R35 GM118096 to M.P. AEB was supported by NIH T32 GM008570. K.Z.P.-V. has been supported by the National Institutes of Health F31 GM131521 and T32 GM007092, and by a Graduate Diversity Enrichment Program Award from the Burroughs Wellcome Fund.

## Author contributions

K.Z Perez-Vale and M. Peifer conceived the study, K. Yow assisted with analysis of *cnoΔRA,* A.E. Byrnes and K.C. Slep solved the structure of Cno’s PDZ domain, and T.M. Finegan quantified cell shapes. All other experiments were carried out by K.Z Perez-Vale. K.Z Perez-Vale, K.C. Slep, and M. Peifer wrote the manuscript with input from the other authors.

## Materials and Methods

### Fly stocks

Fly stocks created and used in this study are listed in Table 2. *yellow white* flies were used as our control and are referred to in the text as wild type, and all the experiments were performed at 25°C. The *cno*ΔRA germline clones were made by heat shocking larvae for 2 hours at 37°C water bath on two consecutive days. After heat shocking, we collected *hsFLP^1^;;P{neoFRT}82B P{ovo−D1−18}3R/ P{neoFRT}82B cno*ΔRA females, in fertile females the germline is homozygous for *cno*ΔRA, and these were then crossed to *cno*ΔRA/TM3, Sb males.

**Table 2:** Fly stocks and antibodies/probes used in this study.

| Fly Stocks |  |  | Source |
| --- | --- | --- | --- |
| *Cno*ΔRA | | | See Materials and Methods |
| *Cno*WT-GFP |  |  | See Materials and Methods |
| *Cno*ΔPDZ-GFP | | | See Materials and Methods |
| *Cno*ΔFAB-GFP | | | See Materials and Methods |
| Antibodies/probes | Specie | Dilution | Source |
| anti-Canoe | Rabbit | 1:1000 (IF, WB) | (Sawyer et al., 2009) |
| anti-Bazooka | Rabbit | 1:2000 (IF) | (Choi et al., 2013) |
| anti-Armadillo (N2 7A1) | Mouse2a | 1:50 (IF),<br>1:500 (WB) | Developmental Studies Hybridoma Bank (University of Iowa, Iowa City, IA, USA) |
| anti- Polychaetoid (PYD2) | Mouse2b | 1:250 (IF),<br>1:500 (WB) | Developmental Studies Hybridoma Bank |
| anti-α-Tubulin (T6199) | Mouse | 1:5000 (WB) | MilliporeSigma (St. Louis, MO) |
| anti-GFP (JL-8) | Mouse2a | 1:1000 (WB) | Clontech Laboratories (Mountain View, CA) |
| anti-GFP (4C9) | MouseG1 | 1:50 (IF) | Developmental Studies Hybridoma Bank |
| anti-GFP (Ab13970) | Chicken | 1:1000 (IF) | Abcam (Cambridge, United Kingdom) |
| Antibodies: |  |  |  |
| Alexa 488, 568, and 647 |  | 1:1000 (IF) | Life Technologies (Carlsbad, CA) |
| IRDye 680DR | Rabbit | 1:15000 (WB) | Licor Biosciences (Lincoln, NE) |
| IRDye 800CW | Mouse | 1:15000 (WB) | Licor Biosciences |

### Generation of *cno*ΔRA allele

The *cno*ΔRA allele was generated using CRISPR/Cas9 to replace part of the *cno* locus, starting 605bp upstream of the translation start site and finishing 1398bp downstream of exon 6, for a total of 3687bp deleted via homology-direct repair (HDR).

#### pU6-gRNAs

The *cno* gene spans ∼47.5Kbp, and it contains two large introns; thus, it was challenging to delete the whole gene. Using the flyCRISPR Target Finder (http://tools.flycrispr.molbio.wisc.edu/targetFinder/) we identified guide RNAs (gRNAs) that would remove a region of the locus that deleted the start codon and multiple exons, reasoning that this would lead to a null allele. We used the maximum stringency on the flyCRISPR Target Finder to identify gRNAs that were 20bp in length with NGG PAM sites only and zero predicted off-target effects. The gRNA regions were sequence-verified to be present in the fly stock to be injected. The sense and antisense gRNAs oligos were annealed and cloned into the pU6-BbsI-chiRNA vector via the BbsI restriction site. Before injection, all the constructs created in this study were verified by PCR, restriction digest, and Sanger sequencing. Below are the gRNA oligonucleotides used.

Oligo nucleotides for gRNA1 (Target (PAM underlined)): GTTTCGATTTATGATCGGTC<u>GGG</u>

- Sense oligo: 5′-CTTCGTTTCGATTTATGATCGGTC-3′
- Antisense oligo: 5′-AAACGACCGATCATAAATCGAAAC-3′

Oligo nucleotides for gRNA2 (Target (PAM underlined)): GTCTCTCTTCAAAGTCCCCT<u>GGG</u>

- Sense oligo: 5′-CTTCGTCTCTCTTCAAAGTCCCCT-3′
- Antisense oligo: 5′-AAACAGGGGACTTTGAAGAGAGAC-3′

#### Donor template

We used the pHD-DsRed-attP as the donor vector, as described in (Gratz et al., 2014). This vector contains an attP phage recombination site and a 3xP3-DsRed cassette (flanked by loxP), which expresses Dsred in the adult eye, which we used as a positive marker to screen for this new allele. We created the dsDNA donor template for HDR using the pHD-DsRed-attP vector by cloning upstream and downstream homology arms flanking the gRNAs adding AarI and SapI restriction sites, respectively. Both upstream and downstream homology arms were 983bp, plus they were flanked by AarI and SapI sites, respectively. These homology arms were PCR amplified (Phusion Polymerase, New England Biolabs) from *yellow white* fly genomic DNA using the primers described below.

Primers for PCR amplification of the upstream homology arm (AarI sites are underlined; **target sequences** are bold):

- Forward: 5′-GACT<u>CACCTGC</u>ATCGTCGC**CGAAACGAAATTTATATTTACCAGC**-3′
- Reverse: 5′-GACT<u>CACCTGC</u>ATCGCTAC**GTCGGGCTAACATATAGCCAATTGA**-3′

Primers for PCR amplification of the downstream homology arm (SapI sites are underlined; **target sequences** are bold):

Forward: 5′-GACT<u>GCTCTTC</u>ATAT**CCTGGGCCATTCGAGAGGTGTA**-3′

Reverse: 5′-GACT<u>GCTCTTC</u>AGAC**GGAACGTGCTACAACCTCAAA**-3′

#### Embryo injections and screening

A mixture containing the two gRNA pU6-BbsI-chiRNA vectors (75 ng/μL per gRNA) and the HDR pHD-DsRed-attP plasmid with the homology arms (250 ng/μL) was sent to BestGene (Chino Hills, CA) for injections. BestGene performed injections of the above mixture into *Cas9(x);;P{ry[+t7.2]=neoFRT}82B ry[506]* embryos. To detect germline transmission of our CRISPR modifications, we outcrossed the G0 adults to *yellow white*. Offspring (F1) were screened for the presence of dsRed eye fluorescence, which indicates there was homology-direct repair (HDR) with our dsDNA donor.

The positive flies were then crossed to *w^-^; TM6B, Tb/TM3, Sb* to generate a balanced stock.

#### Molecular characterization of engineered cno allele

The deletion and the position of the deletion on *cno* locus were verified by PCR (Phusion polymerase, New England Biolabs) using the primers below. The forward primer location was upstream of the upstream homology arm, and the reverse primer was downstream of the downstream homology arm.

Primers for PCR verification of *cno* ∼3.7Kb deletion:

- Forward: 5′-GGTGCTTGATATGGGAACAC-3′
- Reverse: 5′-GGATGGAATGGGTTAAGTCA-3′

`o` Sequence verification primer for *cno*ΔRA locus attP site: 5′-GCTTCGAGCCGATTGTTTAG-3′

Primers for PCR verification of *cno* ∼3.7Kb deletion and DHR: Detection of dsRed:

- Forward: 5′-CGAGGACGTCATCAAGGAGT-3′
- Reverse: 5′-GGTGATGTCCAGCTTGGAGT-3′

Detection of upstream homology and dsRed:

- Forward: 5′-GGTGCTTGATATGGGAACAC-3′
- Reverse: 5′-GGTGATGTCCAGCTTGGAGT-3′

Detection of dsRed and downstream homology:

- Forward: 5′-CGAGGACGTCATCAAGGAGT-3′
- Reverse: 5′-GGATGGAATGGGTTAAGTCA-3′

### Wild-type rescue construct for ΦC31-mediated integration into attP site in *cno*ΔRA allele

The *CnoWT-GFP* rescue construct was generated by assembling three PCR-amplified fragments into a linearized (EcoRI/KpnI digest) pGE-attB-GMR (Huang et al., 2009) vector using GeneArt Seamless PLUS Cloning and Assembly (Invitrogen). Two of the PCR-amplified fragments were generated from genomic DNA— the 5’UTR deleted sequence in the *cno*ΔRAallele with an additional 500bp upstream of this deletion (606bp total) and a shorter 3’UTR (355 bp) from *cno* isoform RE. We generated the GFP-tagged *cno* full-length (6,944bp) from CDS from the vector used to create the UASp.*cno*FL-GFP fly stock (Bonello et al., 2018). The primers used to generate these fragments contained overlaps between each other. The construct was verified by PCR, restriction digest, and Sanger sequencing.

Primers used for the upstream region and 5’UTR (forward primer overlaps with pGE-attB-GMR and the reverse primer overlaps with cnoFL-GFP, both overlaps are underlined, and target sequence are bold): Forward: 5′-<u>gggctccccgggcgcgtactccacg</u>**CCGATCATAAATCGAAACGG**-3′

Reverse: 5′-<u>tatcatgtgacat</u>**ACTGTACAGGTTGGGGAAAG**-3′

Primers used for cnoWT::GFP (forward primer overlap with 5’UTR and reverse primer overlaps with 3’UTR, both overlaps are underlined, and target sequence are bold):

Forward: 5′-<u>caacctgtacagt</u>**ATGTCACATGATAAGAAGATGTTG**-3′ Reverse: 5′-<u>cgatcccttcttc</u>**TTACGTCACGTGGACCGG**-3′

Primers used for the 3’UTR (forward primer overlaps with cnoFL::GFP and the reverse primer overlaps with pGE-attB-GMR, both overlaps are underlined, and target sequence are bold):

Forward: 5′-<u>ccacgtgacgtaa</u>**GAAGAAGGGATCGTTGCTTAATG**-3′ Reverse: 5′-<u>catacattatacgaagttatggtac</u>**CGCCTTGCTTTCGTTGCCATGTC**-3′

### Generation of mutant rescue constructs for ΦC31-mediated integration into *cno*ΔRAallele attP site

The *cnoWT-GFP* rescue construct was used to generate two additional constructs one lacking the PDZ domain (1006-1099aa, 94aa) and the other lacking a conserved region of the FAB domain (1937-2051aa, 115aa) using the Q5 Site-Directed Mutagenesis Kit (New England Biolabs). We generated a 282bp deletion of the PDZ domain to create the *cnoΔPDZ-GFP*, extending from 5′-CCGCAACCGGAATTGCAGCT-3′ to 5’-TGGCCAAGCAGGGAGCCATC-3′. We made a 345bp deletion of a consensus sequence for the F-actin binding domain to create the *cnoΔFAB-GFP*, extending from 5′-CAGGATTTGGACACTATCAG-3′ to 5′- GAGATATAGACGCGGTGCAC-3′. The primers used to generate these constructs are below.

Primers used for generating the PDZ domain deletion: Forward: 5′-TATCACGGGTTGGCTACA-3′ Reverse: 5′-AAGTTTGTTAGACATGGCAC-3′

Primers used for generating the FAB domain deletion: Forward: 5′-AAGGGTGGGCGCGCCGAC-3′

Reverse: 5′-ATGCTCCATACCGTAATTATCCATAATATATTGCTGCTGC-3′

#### Embryo injections and screening

Injection of the *cno*WT-GFP rescue construct and the *cnoΔPDZ-GFP* and *cnoΔFAB-GFP* mutant rescue constructs were performed by BestGene (Chino Hills, CA) into *PhiC31/int^DM.Vas^;;cnoΔRA* embryos. Offspring (F1) were screened by the presence of w^+^ and outcrossed to *w-; TM6B, Tb/TM3, Sb* to generate a balanced stock. We verified the integration of *cno*WT::GFP, *cnoΔPDZ- GFP* and *cnoΔFAB-GFP* by Western blot and PCR amplification.

Primers used for identifying the PDZ domain in *cno* genomic DNA and in our rescue constructs sequence: Forward: 5′-GGAATCTGTGGCAGACGAAC-3′

Reverse: 5′-CACGCTGGATCACAGGACTAG-3′

The primers used for detecting the PDZ deletion are the same as the *cnoWT-GFP* verification. Primers used for detecting the FAB deletion:

Forward: 5′-GGAGGAGCGTGAGAAGGATT-3′ Reverse: 5′-GAACTTCAGGGTCAGCTTGC-3′

### Embryo Fixation and Immunofluorescence

Flies were allowed to lay eggs at 25°C on apple juice agar plates with yeast paste. Embryos were collected, and embryo fixation and staining was performed as in (Bonello et al., 2018).. In summary, embryos were dechorionated in 50% bleach, washed three times in 0.03% Triton X-100 with 68 mM NaCl, and then fixed in 95°C Triton salt solution (0.03% Triton X-100 with 68 mM NaCl, 8 mM EGTA) for 10 seconds. We then added ice-cold triton salt solution and transfer to ice for fast cooling for at least 30 minutes. We then devitellinized the embryos by vigorous shaking in 1:1 heptane:methanol solution. The embryos were then washed three times with 95% methanol/5% EGTA and stored in 95% methanol/5% EGTA for at least 48 hours at -20°C before staining. Before staining, the embryos were washed three times with 0.01% Triton X- 100 in PBS (PBS-T). We then blocked in 1% normal goat serum (NGS) in PBS-T (PNT) for 1 hour, and embryos were incubated in primary antibodies overnight at 4°C or 2-3 hours at room temperature. Once incubation finished, we washed three rimes with PNT and incubated embryos in secondary antibodies overnight at 4°C or 2-3 hours at room temperature. Both primary and secondary antibodies were diluted in PNT, and the dilutions used are listed in Table 2. After the secondary antibody incubation, we washed three times with PNT and stored embryos in 50% glycerol until mounted on glass slides using Aquapolymount (Polysciences, Warrington, PA).

### Image Acquisition and Analysis

Fixed embryos were imaged on a confocal laser-scanning microscope (LSM 880; 40x/NA 1.3 Plan-Apochromat oil objective; Carl Zeiss, Jena Germany). Images were processed and maximum intensity projections were generated using ZEN 2009 software. We used Photoshop (Adobe, San Jose, CA) to adjust input levels and brightness and contrast. Analysis of apical-basal positioning on maximum intensity projections (MIPs) was executed as previously described (Choi et al., 2013) and associated heat maps created using GraphPad Prism 9. Briefly, using Zen 2009 software, the z-stacks were cropped to select a region of interest (ROI) on the *xy*-axis of 250x250 pixels for stacks collected using a digital zoom of 2 or 200x200 pixels for stacks with a 1.6 digital zoom. Using the Zen software the z-stacks ROIs dimensions were modified from *yzx* to *xyz* along the *y*-axis from which MIP were generated. To obtain the data to create heat maps, using ImageJ (National Institutes of Health, Bethesda, MD, USA) the MIPs were rotated 90° counterclockwise and the mean intensity was analyzed using the Plot Profile function.

### Canoe SAJ and TCJ enrichment analysis

Data for analyzing SAJ and TCJ enrichment was obtained from z-stacks taken through the embryo using a digital zoom of 1.6 or 2 and a step size of 0.3 µm. First, the total length of the cells was defined by determining slice position of the apical (top) and basal (bottom) of the cells in the stack. For embryos in mid-late stage 5 the SAJs were determined to be at 21.82% of the total length and the TCJ enrichment was assessed at 33.33% more basal to the SAJs. For embryos in late stage 5, the SAJs correspond to 21.82% of the total length and the TCJ enrichment was assessed at 50% more basal to the SAJs.

The Cno TCJ intensity ratio was measured from MIPs of a 1.2-2.4 µm of the apical adherens junctions region of embryos from stage 7. The MIPs were generated from z-stacks taken through the embryo using a digital zoom of 1.6 or 2 and a step size of 0.3 µm. ImageJ software was used to identify the apical adherens junction region from which z-stack MIPs were generated. The mean intensity of Cno was measured using ImageJ (National Institutes of Health, Bethesda, MD, USA) by creating lines using the line tool (line width of 3 pixels) along the bicellular junctions, avoiding TCJs or multicellular junctions, and next creating short lines at TCJ/multicellular junctions at 300% zoom. For each TCJ or four-way junction, the three or four bicellular junctions in contact with that junction were measured to obtain the mean intensity. A total of ten cells were quantified per embryo and a total of four embryos were assessed from three experiments. The average bicellular junction intensity per cell was calculated. The Canoe TCJ ratio was calculated by dividing the mean intensity of the TCJ by the average of the bicellular junctions. Box and whiskers graphs were made using GraphPad: the box shows the 25^th^-75^th^ percentile, the whiskers are 5^th^-95^th^ percentiles, the horizontal line is the median and the plus sign (+) is the mean. Data statistical analysis was done using GraphPad. Statistical significance was calculated by Welch’s unpaired t-test or Brown Forsythe and Welch ANOVA test.

### Gap analysis

Representative fields of cells, visualized at the level of the apical adherens junctions and measuring 133 x 133 µm, were selected from embryos at stage 7 and stage 8, matching stages for each genotype. Images were visually inspected in the Arm channel, looking for gaps at tricellular junctions/rosette centers or along aligned AP borders that exceeded roughly 1µm. For long gaps along aligned AP borders, one gap was counted per ∼4 cells (roughly the number of cells that would have formed a rosette at that location).

### Planar polarity quantification

The planar polarity of Cno, Baz, Arm, and Pyd was measured from MIP of a 2.4 µm region of the apical adherens junctions of embryos from stage 7 to early stage 8. The MIPs were generated from z-stacks taken through the embryo using a digital zoom of 1.6 or 2 and a step size of 0.3 µm. Using ImageJ software the 2.4 µm region was identified from which MIPs were generated. The mean intensity and orientation (angle) was measured using ImageJ (National Institutes of Health, Bethesda, MD, USA) by creating lines (line width of 3 pixels) at 300% zoom at bicellular borders without including the TCJ/multicellular junctions. Lines at border were sorted by angle relative to the AP axis. AP borders (vertical) were defined as those at 60°-90° and the DV borders (horizontal) at 0°-29°. The background mean intensity for Cno, Baz, Arm and Pyd was measured by drawing circles in the cytoplasmic region of 13 cells, and calculating the average of their mean intensities. This background average was subtracted from the average of the AP and DV border measurements to obtain the average intensity for the AP and DV border. A total of four embryos were assessed from at least three experiments. Baz and Pyd were normalized to AP borders producing a DV/AP ratio, and Cno and Arm were normalized to DV borders producing a AP/DV ratio. Box and whiskers graphs were made using GraphPad.

The box shows the 25^th^-75^th^ percentile, the whiskers show 5^th^-95^th^ percentiles, the horizontal line shows the median and the plus sign (+) is the mean. Data statistical analysis was done using GraphPad. Statistical significance was calculated by Welch’s unpaired t-test or Brown Forsythe and Welch ANOVA test.

### Cell topology analysis

The ventrolateral regions of embryos stained with an AJ marker were extracted and were processed by hand and converted to binary to ensure accurate segmentation. These images were aligned such that the anterior-posterior axes were oriented left-right. Images were then processed by a custom Matlab script to automatically segment cells and the shape characteristics were extracted using the ‘regionprops’ functionality. Data was then analyzed for statistical significance and graphs made using Prism software (GraphPad). Statistical significance was calculated by an unpaired t-test with Welch’s correction.

### Cuticle preparation

Cuticle preparation was performed according to (Wieschaus and Nüsslein-Volhard, 1986). Briefly, embryos were collected and aligned on an apple juice agar plate and incubated at 25°C for 48 hours to allow embryos to develop fully. All unhatched embryos were collected in 0.1% Triton X-100 and dechorionated in 50% bleach for 5 minutes. They were then washed three times with 0.1% Triton X-100 and transferred to glass slides where all the liquid was removed and mounted in 1:1 Hoyer’s medium:lactic acid and incubated at 60°C for 24-48 hours. They were then stored at room temperature.

### Western blotting

Table 2 contains the antibodies and dilutions used for these experiments. Protein levels expression of Cno, Pyd, and Arm were determined by immunoblotting embryos collected in the 1-4 hours and 12-15 hours windows. The lysates were generated as in (Manning et al., 2019). Briefly, embryos were dechorionated for 5 minutes in 50% bleach. After washing three times with 0.1% Triton X-100, lysis buffer (1% NP-40, 0.5% Na deoxycholate, 0.1% SDS, 50 mM Tris pH 8.0, 300 mM NaCl, 1.0 mM DTT, 1x Halt protease, phosphatase inhibitor cocktail (100x), and 1 mM EDTA) was added and the embryos were placed on ice. Embryos were ground in a microcentrifuge tube using a pestle, lysate was centrifugated at 13200 RPM for 15 minutes at 4°C and protein concentration was determine using Bio-Rad Protein Assay Dye. The lysates were resolved using 7% SDS-PAGE and transferred to nitrocellulose membrane. The membranes were incubated with primary antibody either for 2 hours at room temperature or overnight at 4°C. A 45 minute incubation at room temperature was performed for the secondary antibody. The membranes were developed using the Odyssey CLx infrared system (LI-COR Biosciences). Analysis of band densitometry was calculated using LI-COR Image Studio.

### Cloning and purification

DNA encoding the Drosophila Canoe PDZ domain (amino acid residues 1006-1099) was fused to DNA encoding the C-terminal six amino acids of Drosophila Echinoid (Ed) (amino acid sequence IREIIV) using the PCR method (Forward Primer: 5’-GGCAGGACCCATATGccgcaaccggaattgcagctcattaag-3’, Reverse Primer: 5’-GCCGAGCCTGAATTCTTACACAATGATTTCGCGAATgatggctccctgcttggccacttc-3’), and sub-cloned into pET28 (Millipore Sigma, Burlington, MA) using NdeI and EcoRI restriction sites. PDZ-Ed plasmid was transformed into Escherichia coli BL21 DE3 pLysS cells, grown to an optical density at 600 nm of 1.0 in media containing 50 μg/l kanamycin, the temperature lowered to 18° C, and protein expression induced with 100 μM Isopropyl β-D-1-thiogalactopyranoside for 16 hours. Cells were harvested by centrifugation, resuspended in buffer A (25 mM Tris pH 8.0, 200 mM NaCl, 10 mM imidazole, 0.1% β-ME) at 4° C, and lysed by sonication. Phenylmethylsulfonyl fluoride was added to 1 mM final concentration.

Cells debris was pelleted by centrifugation at 23,000 x g for 45 minutes and the supernatant loaded onto a 10 ml Ni2+-NTA column (Qiagen, Hilden, Germany). The column was washed with 600 ml buffer A and protein batch eluted with 100 ml buffer B (buffer B = buffer A supplemented with 290 mM imidazole).

CaCl2 was added to 1 mM final concentration, and 0.1 mg bovine α-thrombin added to proteolytically cleave off the N-terminal His6 tag. After a 20-hour incubation period at 4° C, protein was filtered over 0.5 ml of benzamadine sepharose (Cytiva, Marlborough, MA), concentrated in a Millipore 3k MWCO centrifugal concentrator (Millipore Sigma, Burlington, MA), and diluted into 100 ml buffer C (25 mM HEPES pH 7.0, 0.1% β-ME). PDZ-Ed was loaded onto a 10 ml SP-Sepharose Fast Flow column (Cytiva, Marlborough, MA), washed with 100 ml buffer C, and step eluted with five-50 ml volumes of buffer C supplemented with 100, 200, 300, 400, and 500 mM NaCl respectively. PDZ-Ed-containing fractions were concentrated in a Millipore 3k MWCO centrifugal concentrator (Millipore Sigma, Burlington, MA), exchanged into 25 mM HEPES pH 7.0, 100 mM NaCl, 0.1% β-ME, concentrated to 15 mg/ml, aliquoted, flash frozen in liquid nitrogen, and stored at -80 °C.

### Crystallization, data collection, and structure determination

Canoe PDZ-Ed was crystallized via hanging drop: 2 μl of 15 mg/ml protein plus 2 μl of a 1 ml well solution containing 18% PEG 3350 and 300 mM sodium acetate, 18 °C. PDZ-Ed crystals were frozen in fomblin oil and a native diffraction data set collected on single crystals at the Advanced Photon Source 22-ID beamline at 100 K (400 frames, 0.5° oscillations, 12398.420 eV). Data were processed using HKL2000 (Otwinowski and Minor, 1997). Phasing was obtained via the molecular replacement method (search model: mouse Afadin PDZ, PDB accession code 3AXA, chain A residues 1-91 with the nectin-3 residues removed (Fujiwara et al., 2015)]), yielding a refined log likelihood gain of 691 for two protomers in the asymmetric unit (a.s.u.) and apparent density for the echinoid C-terminal six residues bound to each PDZ domain in the a.s.u.. Initial models were built using AutoBuild (PHENIX) followed by reiterative buildings in Coot (Emsley et al., 2010) and subsequent refinement runs using phenix.refine (PHENIX) (Adams et al., 2010). Refinement runs used real space, simulated annealing refinement protocols (temperatures: 5,000 K start, 300 K final, 50 steps), torsion angle molecular dynamics (temperatures: 2,500 K start, 300 K final, 500 steps), individual B-factor refinement, and atomic displacement parameters using a maximum-likelihood target. The final refinement run produced a Rfree value of 22.6%. The final model includes residues 1011-1096; 1099 (Canoe PDZ) and 1100-1105 (Echinoid C-terminal six residues) for chain A, residues 1008-1099 (Canoe PDZ) and 1100-1105 (Echinoid C-terminal six residues) for chain B, and 112 water molecules. Data collection and refinement statistics are summarized in Table X. Structure images were generated using the PyMOL Molecular Graphics System, version 2.4.0 (Schrödinger, LLC, New York, NY), including use of the program’s structure alignment protocol.

### Data Deposition

Coordinates for the Drosophila melanogaster Canoe PDZ-Echinoid structure have been deposited in the Protein Data Bank under accession code: 7MFW.

---

**Fig S1.**
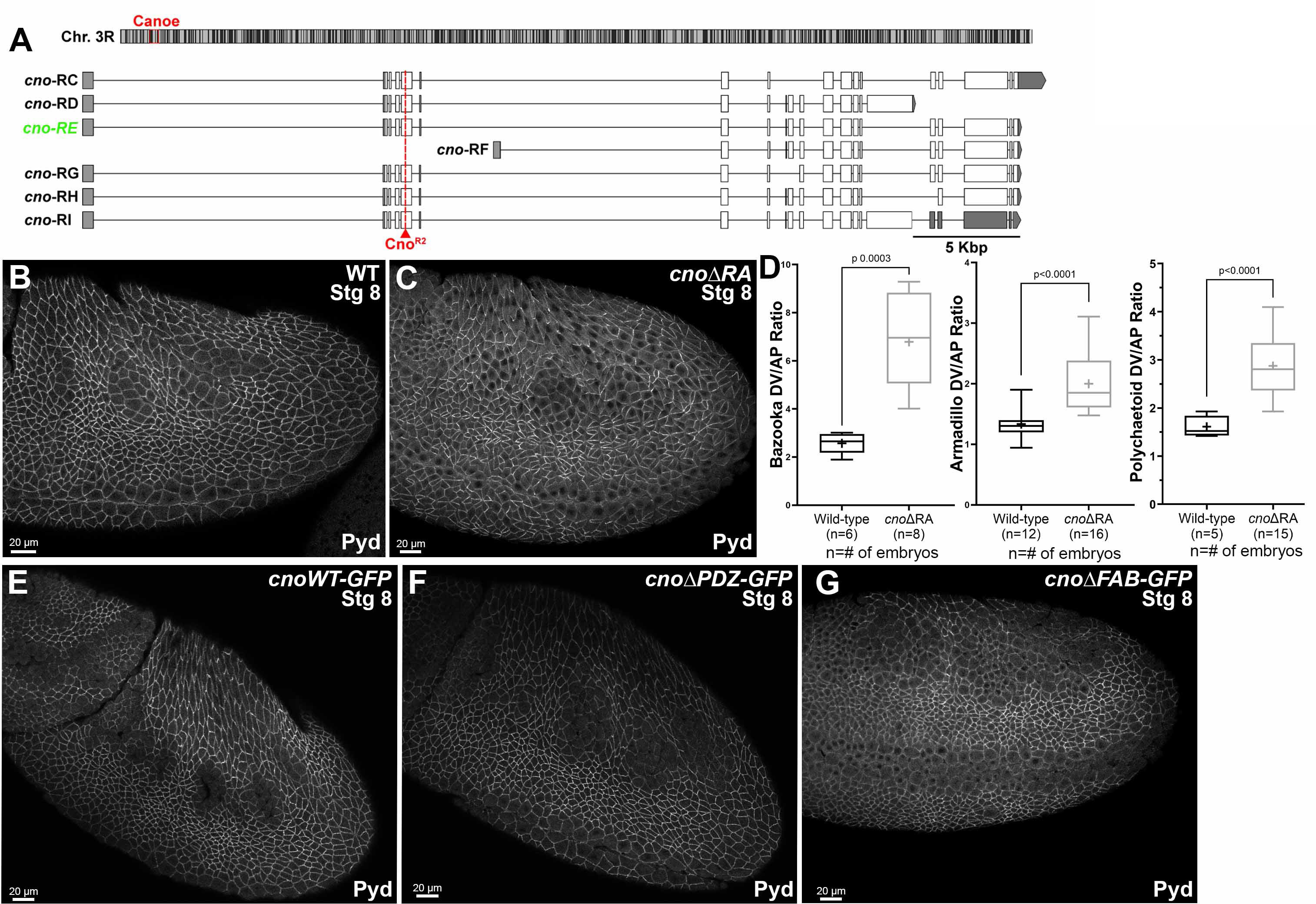
*cno* locus, and aspects of the phenotypes of *cnoΔRA, cnoΔPDZ* and *cnoΔFAB.* A. Top—position of *cno* locus on the third chromosome. Bottom. Predicted mRNAs encoding different *cno* isoforms. Our choice of *cno-RE* was guided by modEncode and other data available on JBrowse at FlyBase. The predicted internal start site of *cno-RF* is not supported by modENCODE or RAMPAGE data (J-Browse), and transcription of its unique first exon is not detected in RNA-seq data until after 20 hours of embryonic development. modENCODE data also suggests the large alternate exon present in *cno-RD* and *cno-RI* is transcribed at lower levels than the other exons. *cno-RE* has the longest coding sequence of the other predicted isoforms. B,C. Stage 8. The ZO-1 homolog Pyd remains localized to AJs in *cnoΔRA* mutants, though like Arm its planar polarization to DV borders is increased. D. Baz, Arm, and Pyd planar polarization are enhanced in *cnoΔRA* mutants. E-G. Stage 8. Pyd localization to AJs appears unchanged in cnoWT, *cnoΔPDZ* and *cnoΔFAB* mutants.

**Fig S2.**
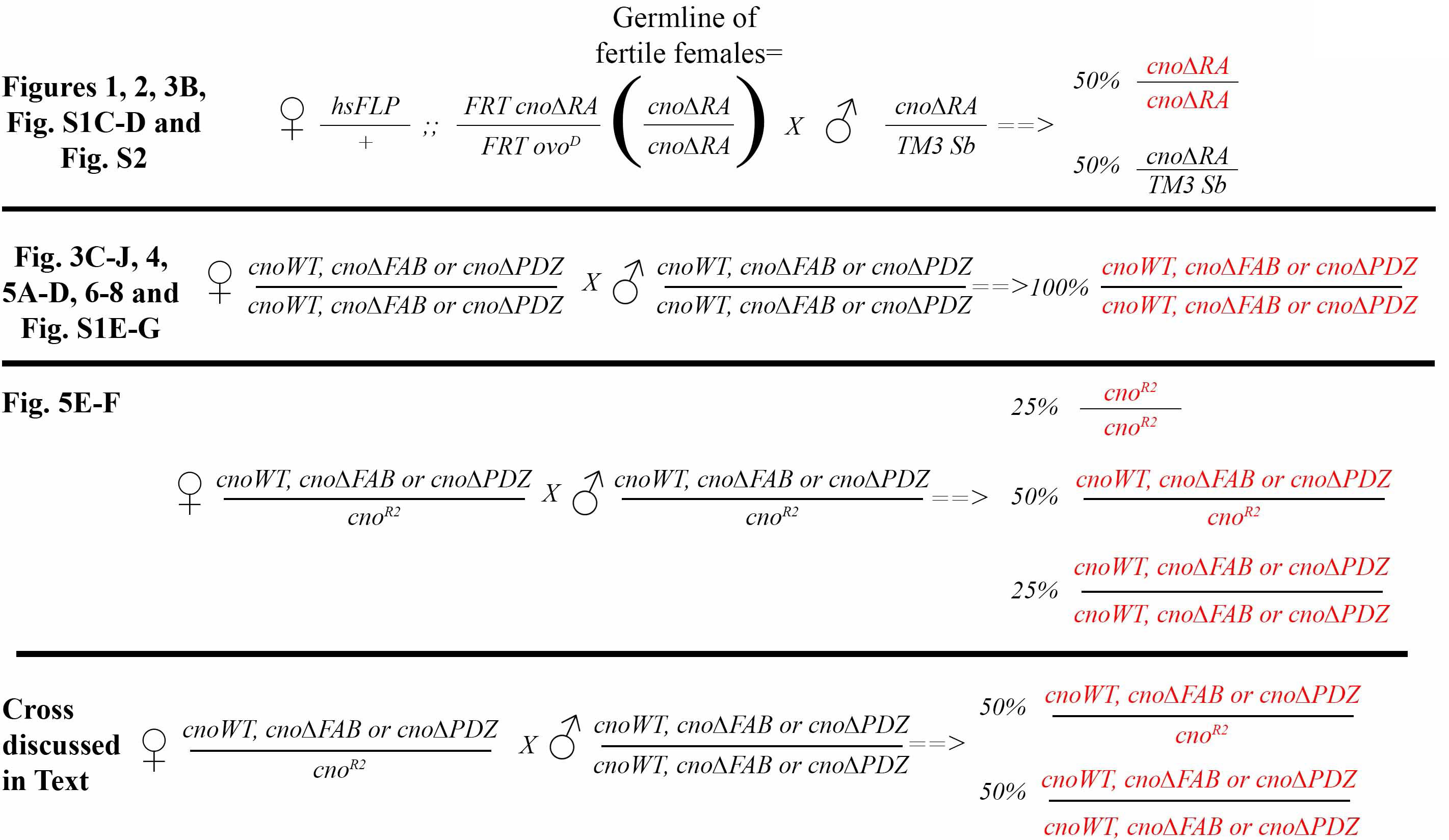
Crosses used to generate embryos analyzed. In the first cross, we used the FLP/FRT approach to mediate site-specific recombination and generate females whose germlines were homozygous for *cnoΔRA* and who thus lost the dominant female sterile mutation *ovo^D^*. Thus the only fertile females contributed only *cnoΔRA* mRNA and protein to their progeny. They were crossed to *cnoΔRA/+* males and thus half the progeny are *cnoΔRA* maternal and zygotic mutants and half receive a paternal wildtype *cno* allele.

**Fig S3.**
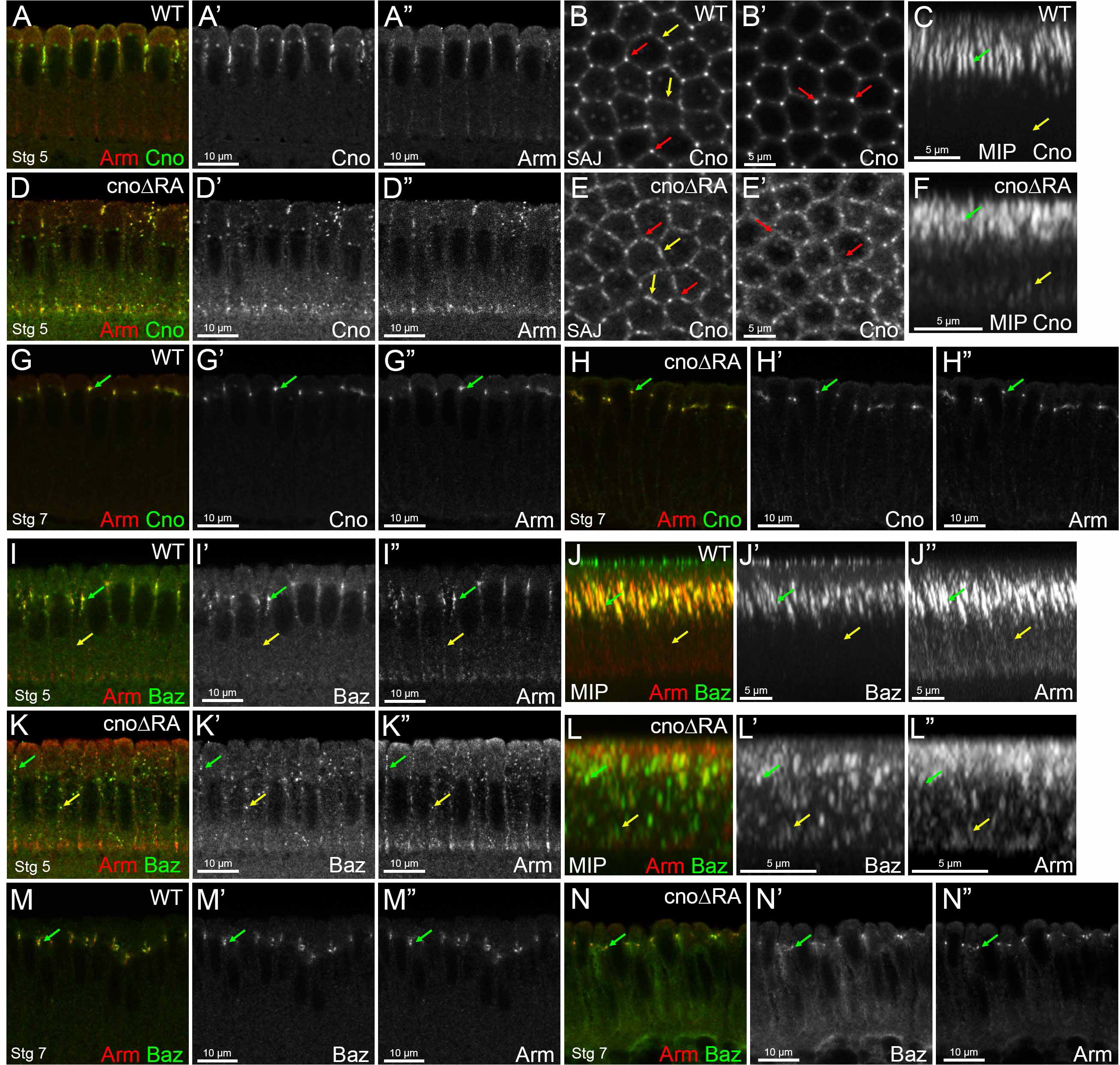
The RA domain is important for Cno localization and its roles in AJ polarization during cellularization. A-H, K-N. Stage 5. I-J,O-P. Stage 7. A,D,G-I,K,M-N. Cross-sections. B,E. *En face* views. C,F,J,L. Maximum intensity projections (MIPs). A-H. Embryos stained with Arm and Cno. I-N. Embryos stained with Arm and Baz. *cno*ΔRA embryos have defects in Cno’s apical restriction (A,A’ vs D,D’) with Cno localizing not only apically but at basal junctions (C vs F, yellow arrows). These mutant embryos have defects in Cno’s TCJ enrichment at SAJs (B vs E, red arrows), localizing more strongly at bicellular junctions (B vs E, yellow arrows). *cno*ΔRA TCJ enrichment defects are even more striking deeper into the embryo (B’ vs E’, red arrows). MIPs reveal that lack of the RA domains affect Cno rod-like structure (C vs F, green arrow). Cno localization is restored during stage 7 in *cno*ΔRA mutant embryos (G vs H, green arrow). *cno*ΔRA embryos have defects in Baz and Arm localization with puncta localizing apically (I vs K, green arrow) and some basolateral (I vs K, yellow arrow). MIP reveal that both Baz and Arm localization is scattered along the membrane (J vs L, yellow arrow) with some apical localization retained (J vs L, green arrow). This localization defect is restored back to WT during stage 7 (M vs N, green arrow).

**Fig S4.**
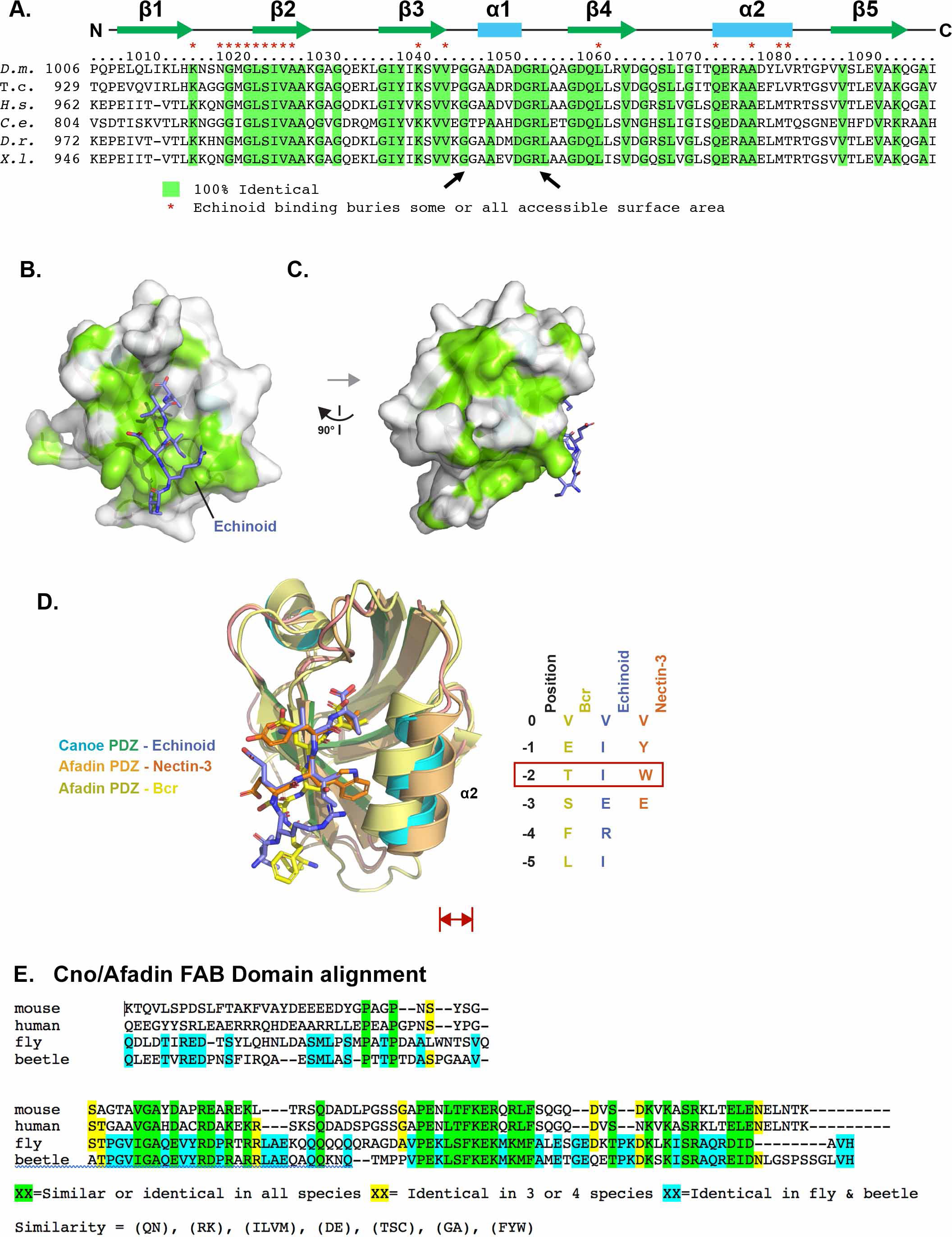
The Cno PDZ domain has extensive, surface exposed conservation, and displays variation in peptide binding across species. A. Alignment of Cno/Afadin homologs from Drosophila melanogaster (D.m.), Tribolium castaneum (T.c.), Homo sapiens (H.s.), Caenorhabditis elegans (C.e.), Danio rerio (D.r.), Xenopus laevis (X.l.). Residues that are 100% identical across all species shown are highlighted in green. Cno PDZ domain residue numbering and secondary structure is depicted at top. Cno PDZ domain residues that become partially or totally buried upon Echinoid binding are denoted above the alignment with a red asterisk. B. Surface structure of the Cno PDZ domain with conservation (as delineated in A) mapped on the surface. The Echinoid C-terminal region is shown in stick format, colored purple. The flexible linker was not resolved in protomer A, which is shown in the Figure, but it was resolved in protomer B, and the Echinoid “peptide” that binds Protomer A is coming in from Protomer B. Pdb code is 7MFW. C. Surface structure of the Cno PDZ domain after a 90° rotation of the structure shown in B. D. Structural alignment of the Cno PDZ:Echinoid structure (this study), the Afadin PDZ-Nectin-3 structure (pdb accession code 3AXA; (Fujiwara et al., 2015), and the AF-6 PDZ-Bcr structure (pdb accession code 2AIN; (Chen et al., 2007). Color coding each PDZ domain and respective bound peptide is indicated. PDZ domains are shown in ribbon format and bound peptides are shown in stick format. Peptide sequences are shown at right, numbered relative to the ultimate valine of each peptide, which is denoted position 0. Residues at position -2 are indicated in a red box, as the size of the side chain at the -2 position correlates with a relative outward shift (red arrows) of the PDZ domain’s α2 helix. E. Sequence alignment of the C-terminal regions of Drosophila (fly) and Tribolium (beetle) Cno and mouse and human Afadin, illustrating the region deleted in CnoΔFAB. The C-terminal 68 amino acids are highly conserved in insects (81% identity) and share conservation in all homologs (41% similarity). More N-terminal is a region conserved only in insects (39% identity), and are not conserved between insects and vertebrates or even between mouse and human Afadin. Sequences immediately more N-terminal are not well conserved among any homologs, and begin to include the homopolymeric runs of amino acids found in the intrinsically disordered domain.

**Fig S5.**
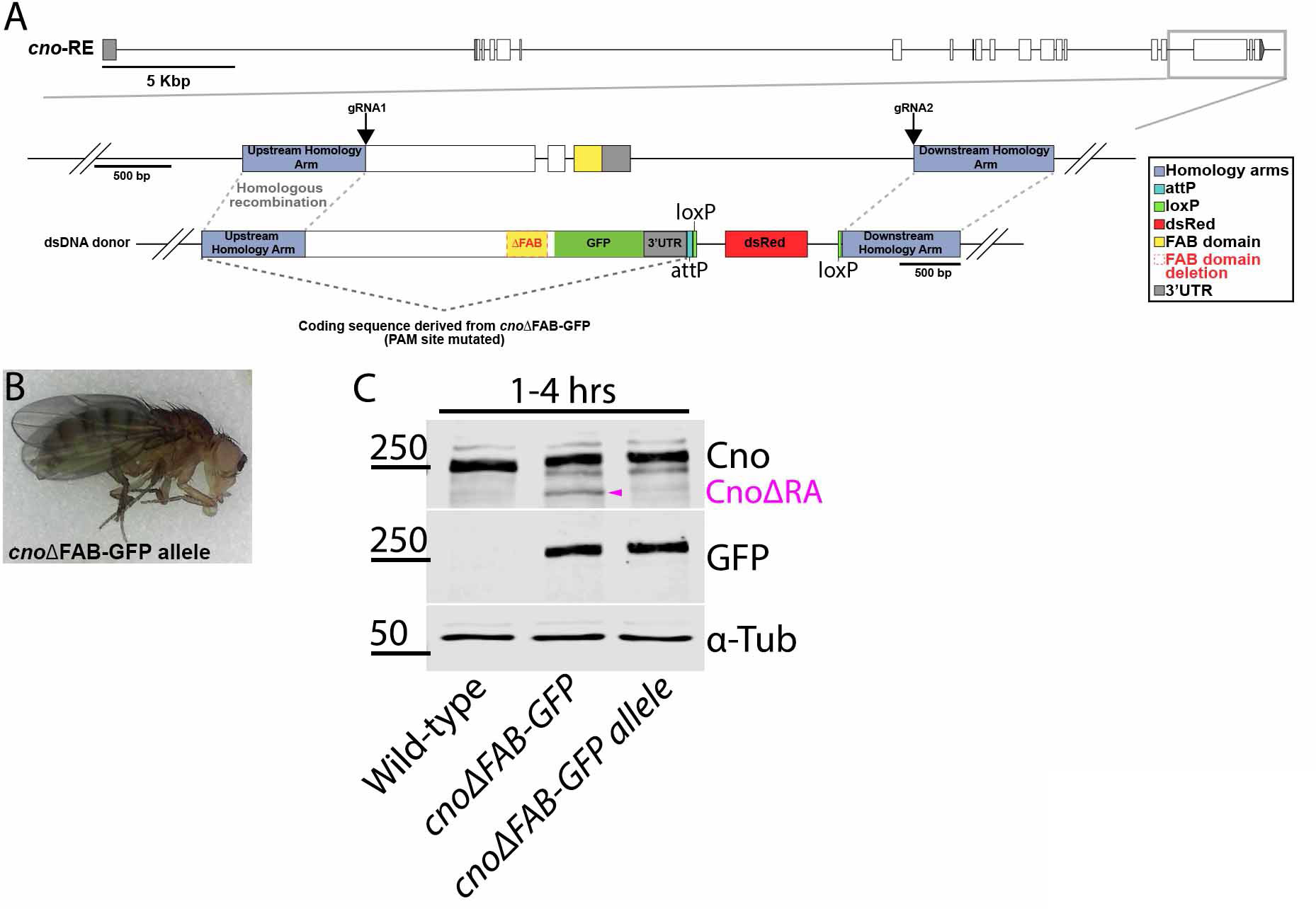
Generating a second *cnoΔFAB* allele directly in the *cno* locus. A. Diagram of the cno genomic locus. Gray box highlights the region manipulated. gRNA 1 target Exon 18 and gRNA 2 targets a sequence downstream of the 3’UTR. dsDNA donor was provided for repair via homologous recombination. The dsDNA donor coding sequence is derived from cnoΔFAB-GFP with the gRNA 1 PAM site mutated. B. Adult fly from stock homozygous for cnoΔFAB-GFP (allele). C. Immunoblot, embryonic extracts from 1-4 hrs, blotted with antibodies to Cno, GFP, and alpha-tubulin as a loading control. Magenta arrow=cnoΔRA.

